# Phospholipid Scramblase 1 (PLSCR1) Regulates Interferon-Lambda Receptor 1 (IFN-λR1) and IFN-λ Signaling in Influenza A Virus (IAV) Infection

**DOI:** 10.1101/2024.11.20.624469

**Authors:** Alina X. Yang, Lisa Ramos-Rodriguez, Parand Sorkhdini, Dongqin Yang, Carmelissa Norbrun, Sonoor Majid, Sanghyun Lee, Yong Zhang, Michael J. Holtzman, David F. Boyd, Yang Zhou

## Abstract

Phospholipid scramblase 1 (PLSCR1) is an antiviral interferon-stimulated gene (ISG) that has several known anti-influenza functions. However, the mechanisms in relation to its expression compartment and enzymatic activity have not been completely explored. Moreover, only limited animal models have been studied to delineate its role at the tissue level in influenza infections. Our results showed that Plscr1 expression was highly induced by influenza A virus (IAV) infection *in vivo* and in airway epithelial cells treated with IFN-λ. We found that infected *Plscr1^-/-^*mice exhibited exacerbated body weight loss, decreased survival rates, heightened viral replication, and increased lung damage. Interestingly, transcriptomic analyses demonstrated that Plscr1 was required for the expression of type 3 interferon receptor (Ifn-λr1) and a large subset of ISGs upon IAV infection. The impaired expression of Ifn-λr1 and downstream ISGs may be responsible for delayed viral clearance and worse lung inflammation in *Plscr1^-/-^* mice. PLSCR1 acts as a transcriptional activator of *IFN-*λ*R1* by directly binding to its promotor after IAV infection. In addition, PLSCR1 interacted with IFN-λR1 on the cell surface of pulmonary epithelial cells following IAV infection, suggesting it also modulated IFN-λ signaling via protein-protein interactions. The lipid scramblase activity of PLSCR1 was found to be dispensable for its anti-flu activity. Finally, single-cell RNA sequencing data indicated that *Plscr1* expression was significantly upregulated in ciliated airway epithelial cells in mice following IAV infection. Consistently, *Plscr1^floxStop^Foxj1-Cre^+^* mice with ciliated epithelial cell-specific Plscr1 overexpression showed reduced susceptibility to IAV infection, less inflammation and enhanced Ifn-λr1 expression, suggesting that Plscr1 primarily regulates type 3 IFN signaling as a cell intrinsic defense factor against IAV in ciliated airway epithelial cells. Our research will elucidate virus-host interactions and pave the way for the development of novel anti-influenza drugs that target human elements like PLSCR1, thereby mitigating the emergence of drug-resistant IAV strains.

## INTRODUCTION

Influenza A virus (IAV) is highly contagious and causes acute respiratory infectious disease transmitted by virus-containing droplets. Its segmented gene and wide host range facilitate frequent antigenic shift during coinfection, posing a significant threat to human health. IAV has caused several pandemics in human history, and seasonal flu remains a major health burden in present days, despite annual vaccination efforts[1]. Existing anti-flu drugs mainly target influenza virus proteins: 1) penetration and shelling inhibitors, such as Gocovri (amantadine) and Flumadine (Rimantadine); 2) neuraminidase inhibitors, such as Relenza (zanamivir), Tamiflu (oseltamivir phosphate), Rapivab (peramivir) and Xofluza (baloxavir marboxil)[2]. However, the emergence of drug-resistant variant strains is common. Moreover, these direct antivirals have a short therapeutic window and are the most effective only if given within the first 48 hours after the initial infection[2]. In addition, IAV-induced acute inflammation could persist, leading to severe complications such as life-threatening pneumonia, immunopathology, and acute respiratory distress syndrome (ARDS). Therefore, there is a pressing need to understand host immune responses and develop anti-influenza drugs targeting host factors.

At the center of anti-flu immunity are the interferon (IFN) pathways. Discovered a mere two decades ago, type 3 IFNs were initially perceived as redundant to type 1 IFNs, given their shared intracellular signaling pathways and antiviral activities. However, recent studies have revealed their unique properties, notably their signaling through a pair of heterodimer receptors (IFN-λR1/IL-10R2) distinct from type 1 IFN receptor complex (IFN-αR1/IFN-αR2)[3, 4]. First of all, IFN-λ exhibits a more constrained expression pattern compared to type 1 IFNs. While the cellular sources of type 1 IFNs in viral infections depend on the infection route and the tissue tropism, they can be produced by a large variety of cell types, including epithelial, parenchymal, immune and stromal cells, to combat infections in the skin, mucosal, organ, lymph node, and at the systemic level[5]. In contrast, IFN-λ is primarily secreted by cells at barrier surfaces, such as respiratory and gastrointestinal epithelial cells, DCs and macrophages[6, 7]. Moreover, their receptor distributions are vastly different. IFN-αR1/IFN-αR2 complex is present on nearly all cell types, but IFN-λR1/IL-10R2 complex is expressed exclusively on epithelial cells and few immune cells, including neutrophils and subsets of dendritic cells[8, 9]. Most importantly, IFN-λ is produced earlier than type 1 interferons in IAV infection and can elicit effective antiviral responses without inflammation such as the release of tissue-damaging mediators such as tumor necrosis factor (TNF) and IL-1β from neutrophils. In fact, only under a high dose of IAV are type 1 interferons detected, contributing to tissue immunopathology [10]. Another exclusive anti-flu mechanism of IFN-λ is its role in preventing viral transmission from the upper airways to the lungs[11]. Taken together, IFN-λ provides a non-redundant front-line shield against influenza virus.

IFNs secreted by infected cells signal neighboring cells to enter an antiviral state by inducing the expression of hundreds of ISGs[12]. PLSCR1, the most studied member of the phospholipid scramblase protein family, is one such ISG, with its expression highly induced by type 1, 2 and 3 interferons in various viral infections[13–15]. It is a type II transmembrane protein generally located on the cell membrane, but can be imported into the nucleus and act as a transcriptional factor to regulate several gene expressions by directly binding to their promoter regions[16, 17]. PLSCR1 is detected in all tissues and has a relatively high expression in lung. Although it is universally expressed across various cell types in the lung including alveolar epithelial cells and lymphocytes, macrophages have the highest expression, followed by endothelial cells and respiratory ciliated cells[18]. The main function of PLSCR1 is to catalyze Ca^2+^-dependent, ATP-independent, bidirectional and non-specific translocation of phospholipids between inner & outer leaflet of plasma membrane. Scrambling of membrane phospholipids results in the externalization of phosphatidylserine (PS), which acts as a docking site for many biological processes including coagulation, apoptosis, and activation[19].

Previous studies have described some critical anti-influenza activities of PLSCR1. For example, PLSCR1 interacts with the nucleoprotein (NP) of IAV, impairing its nuclear import and thereby suppressing virus replication in A549 cells[20]. In addition, Plscr1 competes with immunoglobulin-like domain-containing receptor 1 (ILDR1) for NP binding, inhibiting swine influenza virus (SIV) infection in mice[21]. Besides direct interactions with the virus, PLSCR1 interacts with toll-like receptor (TLR) 9 and regulates its trafficking and ability to induce type I interferon production in plasmacytoid dendritic cells[22]. Furthermore, PLSCR1 is required to potentiate the expression of numerous other ISGs in response to IFN-β in Hey1B cells, including p56, ISG15 and OAS[23]. The goals of this project are to determine the roles of Plscr1 in an IAV-infected mouse model, to implicate its involvement in IFN-λ signaling, and to elucidate the cell types responsible for Plscr1-mediated anti-influenza activities.

To address this knowledge gap, we have employed a systematic approach, leveraging both global and cell type-specific *Plscr1^-/-^* and *Plscr1* knock-in overexpression mice infected with mouse-adapted human IAV. Human respiratory epithelial cell lines harboring PLSCR1 mutations have also been used to elucidate the subcellular roles of PLSCR1. Our findings demonstrate that Plscr1 is an IFN-λ-stimulated gene, and it protects mice against IAV Infection by regulating *IFN-*λ*R1* gene expression in the nucleus and interacting with IFN-λR1 protein on the cell membrane in ciliated airway epithelial cells. Our research will aid in the understanding of virus-host interactions and the development of novel anti-influenza therapeutics targeting human components, preventing the emergence of drug-resistant IAV strains.

## RESULTS

### Plscr1 Suppresses Viral Replication and Protects Mice in IAV Infection

To uncover any anti-influenza functions of Plscr1 in mice, we employed *Wt* and *Plscr1^-/-^*mice and exposed them to IAV infection. We pursued a systematic approach including multiple endpoints and parameters to monitor mouse health, viral replication and dissemination, antiviral and inflammatory responses, and tissue damage (Figure 1A). We established 300 plaque-forming unit (pfu) of IAV as the sublethal dose, and 900 pfu as the lethal dose resulting in ∼50% mortality in *Wt* mice (LD50). Significantly higher *Plscr1* expression was observed in *Wt* infected mice at 3 days post infection (dpi) (Figure 1B), suggesting that Plscr1 is induced and potentially functions as an antiviral ISG in IAV infection *in vivo*.

**Figure 1.**
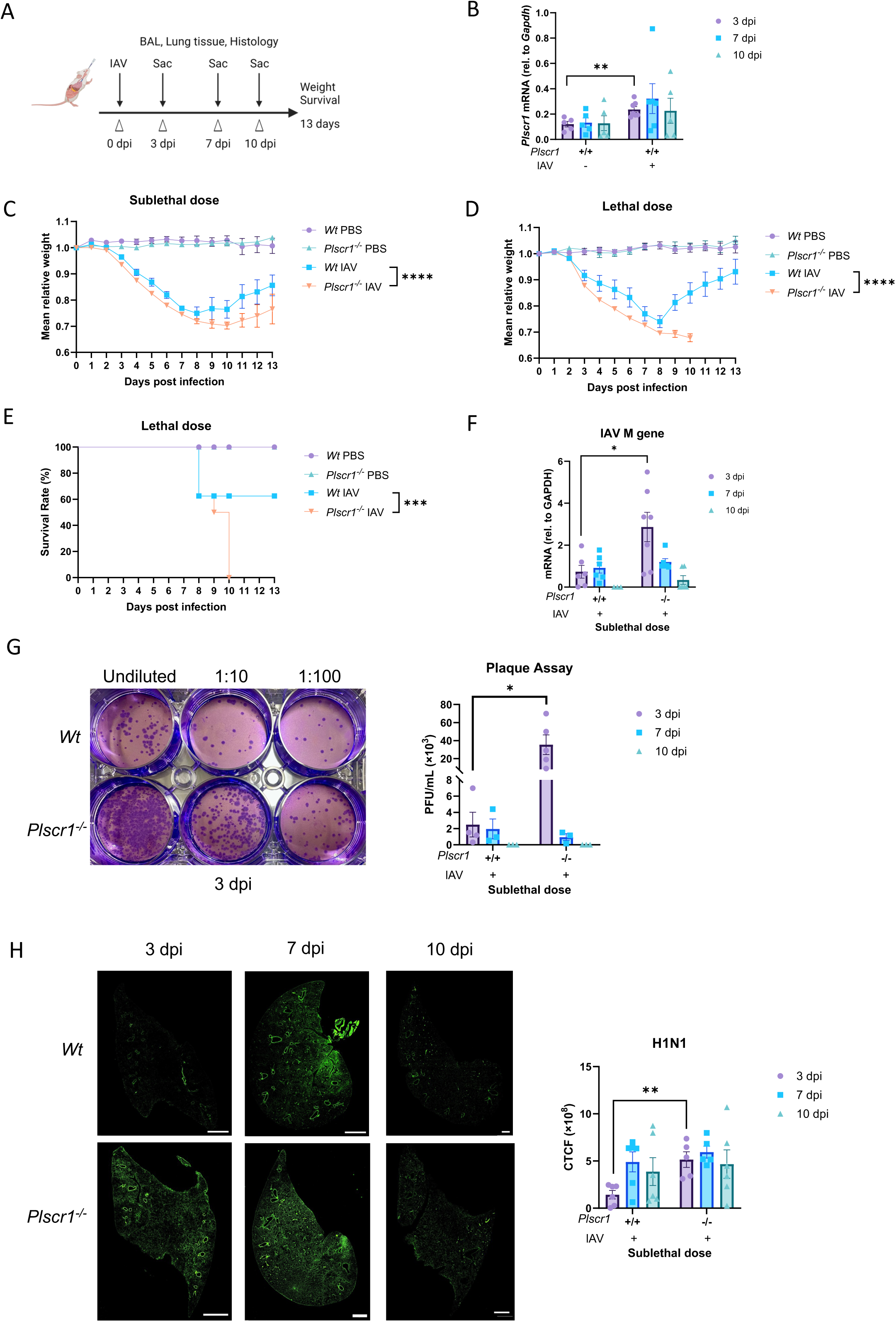
Increased Susceptibility of *Plscr1^-/-^* Mice to Influenza Virus Infection. *Wt* and *Plscr1^-/-^* mice were exposed to sublethal (300 pfu, B, C and F-H) or lethal (900 pfu, D and E) IAV (WSN) infection. (A) Scheme of experiment. (B) Whole lungs of *Wt* mice were analyzed for *Plscr1* RNA by qRT-PCR. (C and D) Mean relative weight of mice post sublethal or lethal infection. (E) Survival rate of mice post lethal IAV infection. (F) Viral RNA load in the lungs was assessed by quantifying M gene by qRT-PCR. (G) Infectious viral titer in the lungs was assessed by plaque assays. (H) Representative staining for H1N1 in lungs. The scale bars represent 1 mm. Quantification was performed using ImageJ. Data are expressed as mean ± SEM of n = 30 mice/group for weight loss post sublethal infection and n=8 mice/group for weight loss and survival rate post lethal infection. For the rest analysis, n= 5-10 mice/group. All data were pooled from three independent experiments. Logrank (Mantel-Cox) test was used to compare survival rates. Ordinary two-way ANOVA tests were used to compare weight losses. *p < 0.05, **p < 0.01, ***p<0.001, ****p<0.0001. dpi, days post infection. CTCF, Corrected Total Cell Fluorescence.

Weight loss generally started at 3 dpi for both mouse strains. *Wt* infected mice reached their lowest weights around 8 dpi, while *Plscr1^-/-^*infected mice experienced continued weight loss for an additional 2 days. *Plscr1^-/-^*mice exhibited significantly greater weight loss compared to *Wt* mice with both sublethal and lethal dose infection at multiple timepoints (Figure 1C and 1D). Moreover, upon infection with 900pfu lethal dose of IAV, *Plscr1^-/-^* mice had a significantly lower survival rate (Figure 1E). In fact, 100% of *Plscr1^-/-^* mice died with 900 pfu of IAV in our experiment by 10 dpi.

We used three assays to robustly and comprehensively measure the viral burden in these animals. First, qRT-PCR analysis using cryopreserved lungs showed that IAV M gene segment mRNA was significantly higher in *Plscr1^-/-^* lungs at 3 dpi (Figure 1F). This observation was further supported by the plaque assay, which measures infectious virus particles, and the IAV titer was significantly higher in *Plscr1^-/-^*lungs at 3 dpi (Figure 1G). Finally, to visualize IAV burden and spread from upper respiratory tract, paraffin-embedded lung sections were stained for H1N1-FITC. In contrast to *Wt* lungs where IAV infection was mostly local and restricted to major airways, *Plscr1^-/-^*lungs had viral dissemination extending from bronchioles to alveoli, with ∼50% of lungs infected. Moreover, *Plscr1^-/-^* lungs had significantly higher corrected total cell fluorescence (CTCF) than *Wt* lungs at 3 dpi, suggesting an overall higher viral burden during acute infection in the absence of Plscr1 (Figure 1H).

### Plscr1 Limits Innate Immunity-Mediated Inflammation and Lung Damage in IAV Infection

Bronchoalveolar lavage (BAL) was harvested for inflammatory cell counts using Cytospin followed by Diff-Quik stain. Immune cells infiltrated the lungs starting at 3 dpi and persisted at least until 10 dpi. *Plscr1^-/-^*mice had significantly higher total BAL cell counts at 7 dpi compared to *Wt* mice, indicating a heightened inflammatory environment in alveoli during acute infection in the absence of Plscr1 (Figure 2A). IAV attracted neutrophils in early infection (3-7 dpi) and lymphocytes in late infection (7-10 dpi). Interestingly, *Plscr1^-/-^* lungs were significantly more neutrophilic at 3 dpi, and neutrophil populations persisted until 10 dpi (Figure 2B).

**Figure 2.**
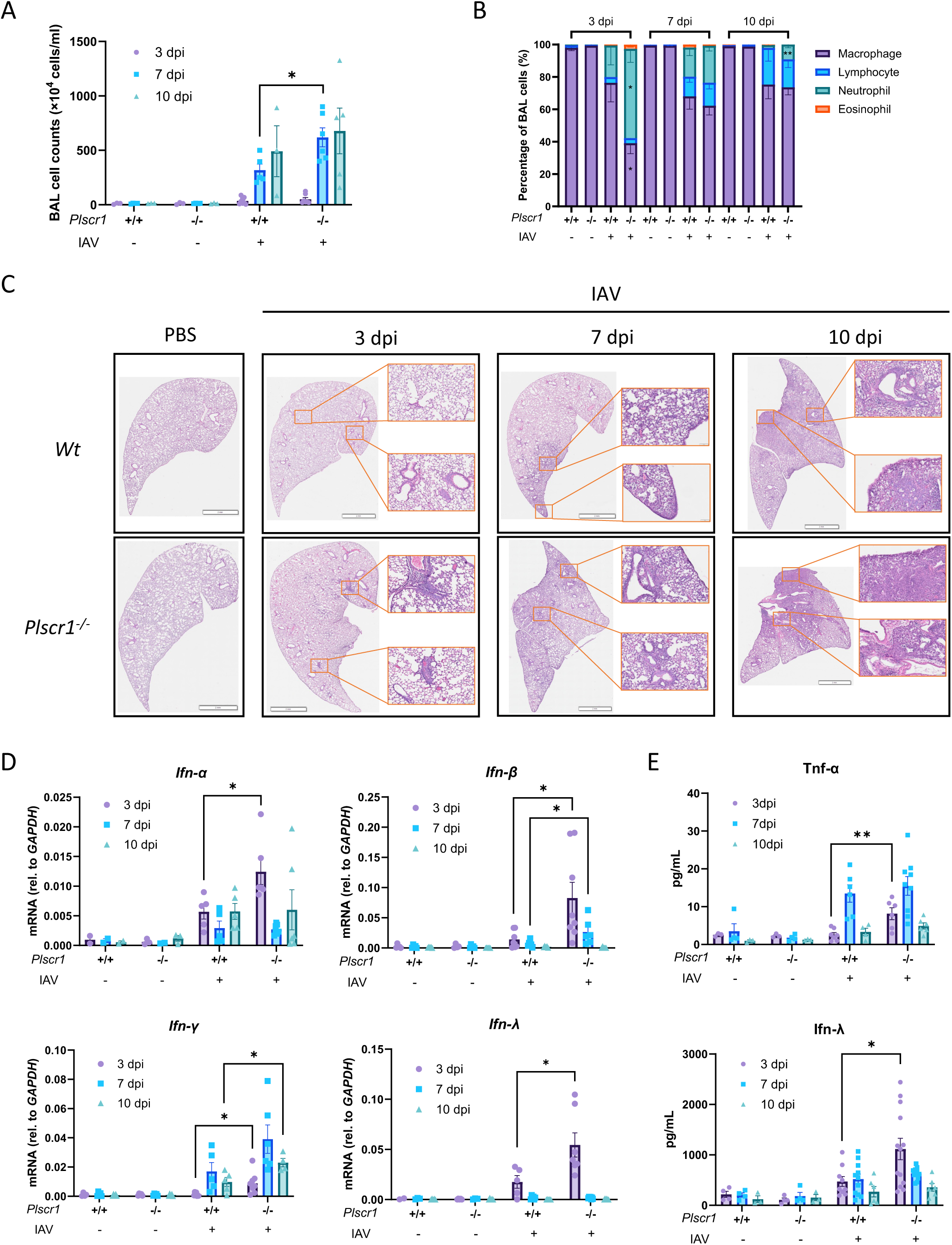
Increased Lung Inflammation in *Plscr1^-/-^* Mice in Influenza Virus Infection. *Wt* and *Plscr1^-/-^* mice were exposed to sublethal (300 pfu) IAV (WSN) infection. (A) Total BAL leukocyte numbers. (B) Differential cell counts in BAL. (C) Representative lung sections stained with H&E. Scale bars represent 3 mm (main) and 200 μm (inlays). (D) Whole lungs were analyzed for *Ifn-*α, *Ifn-*β, *Ifn-*γ and *Ifn-*λ RNA by qRT-PCR. (E) Tnf-α and Ifn-λ concentrations in BAL by ELISA. Data are expressed as mean ± SEM of n = 3-14 mice/group. All data were pooled from three independent experiments. *p < 0.05, **p < 0.01. dpi, days post infection.

Lung sections were stained with Hematoxylin and Eosin (H&E) and histopathology was assessed. *Plscr1*^-/-^ lungs showed minor and localized tissue damage as early as at 3 dpi, when *Wt* lungs appeared normal. Consistently, aggravated immunopathology was evident in *Plscr1^-/-^*lungs at 7 and 10 dpi compared to *Wt* lungs (Figure 2C). These observations included increased thickening and collapse of alveolar walls, pulmonary edema surrounding alveolar walls, inflammatory cell infiltration in peri-bronchial and parenchymal areas, and hyperemia in the absence of Plscr1, suggesting its role in maintaining tissue homeostasis in IAV infection.

To assess the role of Plscr1 as an ISG, IFN expressions in whole lungs were measured by qRT-PCR. We found significantly elevated levels of *Ifn-*α, β, γ and λ expression in *Plscr1^-/-^* mice, particularly in early infection (Figure 2D). This upregulation was associated with heightened production of Tnf-α and Ifn-λ in the BAL at 3 dpi, as assessed by ELISA (Figure 2E). These cytokines may implicate in hyperactive feedforward inflammatory circuits during early IAV infection leading to acute lung injury in *Plscr1^-/-^* mice[24], consistent with worsened histopathology. On the contrary, the expressions of these mediators in *Wt* mice were well controlled, facilitating virus elimination without inciting excessive inflammation. These findings underscore the antiviral and potentially immunoregulatory role of Plscr1.

### Plscr1 Binds to *Ifn-***λ***r1* Promoter and Activates *Ifn-***λ***r1* Transcription in IAV Infection

To determine if IFN signaling pathways are regulated by Plscr1 during IAV infection, RNA sequencing of a total of 20,700 genes was performed using pooled samples from *Wt* and *Plscr1^-/-^* mouse lungs infected with 300 pfu of IAV. A comprehensive examination of interferons and their receptors revealed that *Ifn-*λ*r1* expression was significantly lower in *Plscr1^-/-^*mice, despite high expression of *Ifn-*λ at both 3 and 7 dpi (Figure 3A). Importantly, *Ifn-*λ*r1* was the only IFN receptor exhibiting this pattern, as the expression of its coreceptor *Il10-r*β and other IFN receptors remained unaltered. Inflammatory cytokines including type 1 and 2 IFNs, *Tnf-*α and *Il-1b* were also highly expressed in *Plscr1^-/-^*infected mice, suggesting upstream cytokine production pathways were intact. The RNA sequencing result was further validated by qRT-PCR, which showed that *Plscr1^-/-^* mice failed to upregulate *Ifn-*λ*r1* expression at both 3 and 7 dpi compared to *Wt* mice (Figure 3B). Furthermore, using the online Interferome database[25], we identified a total of 1,113 ISGs in our dataset with a fold change ≥2 (Figure 3C). Enlarged heatmaps with gene names are provided in Supplemental Figure 1. Among those ISGs, 584 are regulated exclusively by type 1 IFNs, and 488 are regulated by both type 1 and type 2 interferons. Unfortunately, the Interferome database does not include information on type 3 IFN-inducible genes in mice. Although many ISGs were robustly upregulated in *Plscr1^-/-^* infected lungs, consistent with inflammation data, a large subset of ISGs failed to be transcribed when *Ifn-*λ*r1* function was impaired, especially at 7 dpi. We suspect that those non-transcribed ISGs in *Plscr1^-/-^* mice may be specifically regulated by type 3 IFN and represent interesting targets for future research. As a result, disrupted type 3 interferon signaling due to impaired expression of *Ifn-*λ*r1* may underlie the delayed viral clearance and exacerbated immunopathology observed in *Plscr1^-/-^*mice[10].

**Figure 3.**
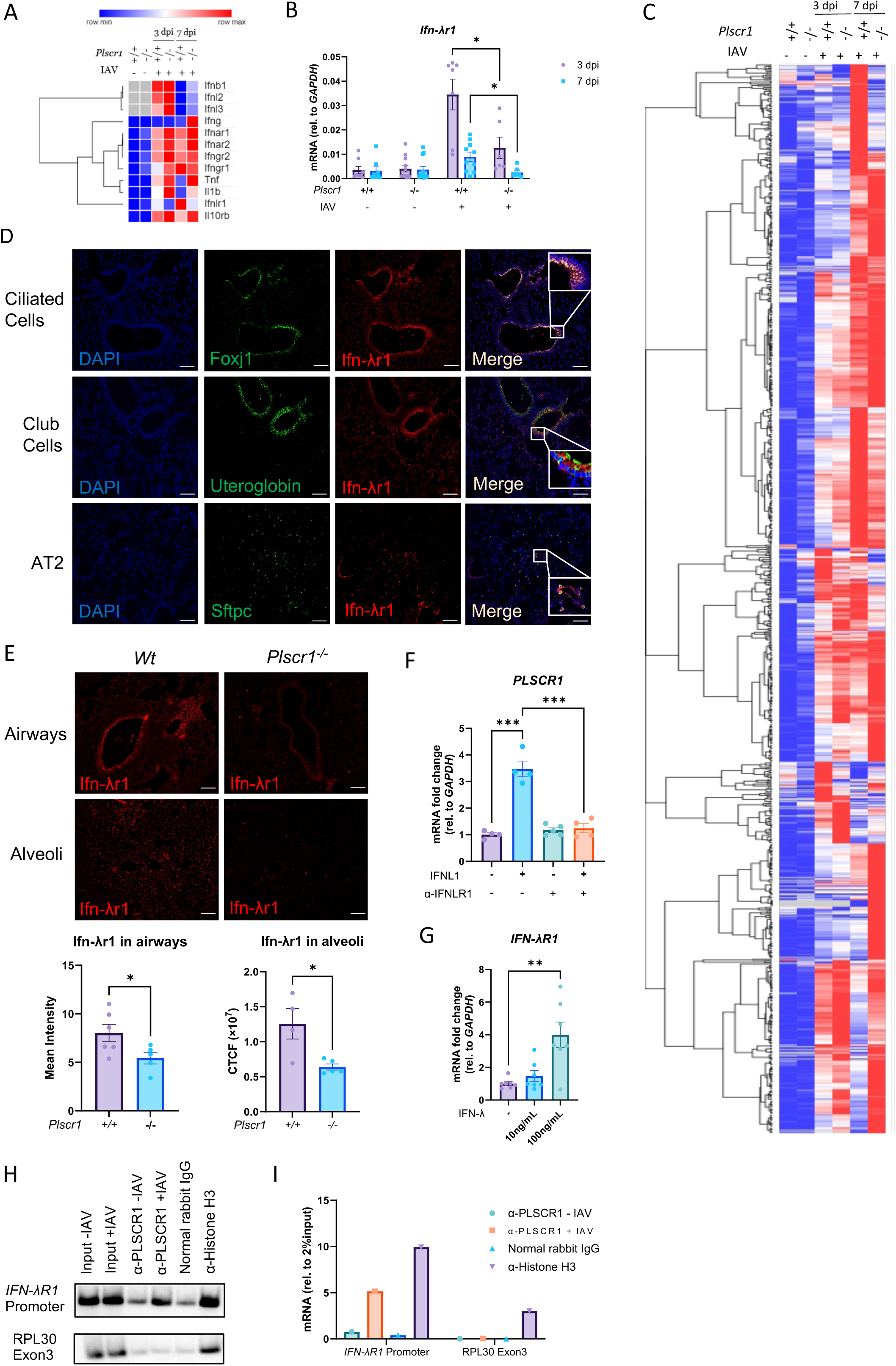
Transcriptional Regulation of *IFN-*λ*R1* by *PLSCR1* and IFN-λ in IAV Infection. (A-E) *Wt* and *Plscr1^-/-^* mice were exposed to sublethal (300 pfu) IAV (WSN) infection. (A) Heatmap of interferons and their receptors in whole lungs by RNA-seq. (B) Whole lungs were analyzed for *Ifn-*λ*r1* by qRT-PCR. (C) Heatmap of differential expressions of all ISGs in whole lungs by RNA-seq. Gene expressions were compared between groups within each row and color-labeled from row minimum (blue) to row maximum (red). (D) Localization of Ifn-λr1+ cells in the lungs of IAV-infected *Wt* mice at 7 dpi. Sections stained for Ifn-λr1 (red), Foxj1, uteroglobin or Sftpc (green) and DAPI (blue) are shown. (E) Representative staining for Ifn-λr1 in airways or alveoli of IAV-infected *Wt* and *Plscr1^-/-^* mice at 7dpi. Quantifications were performed using ImageJ. (F-G) Calu-3 cells were analyzed for *PLSCR1* (F) and *IFN-*λ*R1* (G) RNA by qRT-PCR after recombinant IFN-λ and/or α-IFN-λR1 antibody treatment. Data are presented as fold change compared to non-treated group. (H-I) Chromatin-Immunoprecipitation of PLSCR1 and *IFN-*λ*R1* promoter in Calu-3 cells followed by standard PCR (H) and real-time quantitative PCR (I). Data are expressed as mean ± SEM of n = 4-12 mice or wells/group. For transcriptomic analysis, 9 mice from each PBS-treated group and 4 mice from each IAV-infected group were pooled together. All data were pooled from three independent experiments. *p < 0.05, **p < 0.01, ***p<0.001. dpi, days post infection. CTCF, Corrected Total Cell Fluorescence. Scale bars represent 50 μm.

To corroborate the cell-specific localization of Ifn-λr1 expression, *Wt* IAV-infected mouse lungs were probed with immunofluorescent antibodies to detect Ifn-λr1 and Foxj1 (a marker for ciliated epithelial cells), Uteroglobin (a marker for club cells) or Sftpc (a marker for type 2 alveolar epithelial (AT2) cells). Consistent with previous studies, Ifn-λr1 was present on all of these epithelial cell types (Figure 3D). We further quantified Ifn-λr1 expression in selected airways by measuring mean intensity to disregard the differences in airway areas. Ifn-λr1 expression in alveoli were measured using Corrected Total Cell Fluorescence (CTCF), so that all fluorescent signals within the same size of alveolar areas were analyzed. Expression of Ifn-λr1 was significantly reduced in *Plscr1^-/-^* mouse lungs, in both airways and alveoli regions (Figure 3E). Calu-3 epithelial cells were then used to investigate if *PLSCR1* expression can be regulated by IFN-λ. Using qRT-PCR, we found *PLSCR1* transcription was significantly increased when stimulated with IFN-λ, and this effect was attenuated by pre-incubation with α-hIFNLR1 neutralizing antibody (Figure 3F). Additionally, IFN-λ could directly stimulate the expression of its receptor, *IFN-*λ*R1*, thereby establishing a positive feedback loop to further drive the expression of *PLSCR1* (Figure 3G). These studies demonstrate that *PLSCR1* is an ISG that can be directly stimulated by IFN-λ in airway epithelial cells.

We subsequently investigated the mechanism underlying Plscr1’s transcriptional regulation of *Ifn-*λ*r1* in airway epithelial cells. Chromatin-immunoprecipitation (ChIP) followed by standard PCR in IAV-infected Calu-3 cells unveiled that Plscr1 physically bound to the promoter region of *Ifn-*λ*r1*, with this binding becoming more evident in IAV-infected cells (Figure 3H). These results were further validated using real-time quantitative PCR, suggesting that Plscr1 translocated into the nucleus of lung epithelial cells upon IAV infection to activate *Ifn-*λ*r1* transcription (Figure 3I).

To further determine the regulation of Ifn-λr1 by Plscr1 in response to Ifn-λ signaling without the complexities associated with live virus infection, high molecular weight poly(I:C) was administered intranasally to mice daily for 6 days at 2.5mg/kg of body weight, and mice were sacrificed on the following day (Supplemental Figure 2A). We observed an increase in total BAL cell counts with poly(I:C) administration, although there was no difference between *Wt* and *Plscr1^-/-^*mice (Supplemental Figure 2B). However, *Plscr1^-/-^* mice exhibited a significantly higher number of lymphocytes in BAL (Supplemental Figure 2C), indicating that repetitive poly(I:C) administration might activate adaptive immune responses rather than the innate immune system in an acute IAV infection (Figure 2B). All interferon expressions were elevated following poly(I:C) administration, with comparable levels observed between *Wt* and *Plscr1^-/-^* mice (Supplemental Figure 2D). Consistently, lung histopathology exhibited similar levels of inflammation or tissue damage in both *Wt* and *Plscr1^-/-^*mice (Supplemental Figure 2E). Importantly, *Plscr1* expression was significantly induced by poly(I:C) (Supplemental Figure 2F). Notably, although no other phenotypical or genotypical variances were observed in poly(I:C)-treated *Plscr1^-/-^* mice, *Ifn-*λ*r1* expression was significantly lower in these mice, further affirming the requirement of Plscr1 for *Ifn-*λ*r1* expression in response to IFN-λ (Supplemental Figure 2G).

### Plscr1 Interacts with Ifn-**λ**r1 on Pulmonary Epithelial Cell Membrane in IAV Infection

We employed a combination of multiple biophysical approaches to fully assess the potential protein interaction between Plscr1 and Ifn-λr1. Co-immunoprecipitation (Co-IP) followed by western blot demonstrated that Plscr1 successfully pulled down Ifn-λr1 in both uninfected and IAV-infected *Wt* mouse lungs (Figure 4A and Supplemental Figure 3). Since the co-IP assay may detect indirect interactions through intermediate partners, we used proximity ligation assay (PLA) as a more precise method to identify direct interactions. Consistently, using PLA (Figure 4B), we detected direct interactions between Ifn-λr1 and Plscr1 on airway and alveolar epithelial cells in *Wt* mouse lungs. Noteworthily, significantly more and stronger PLA signals per area were detected in IAV-infected lungs compared to uninfected lungs. In agreement with the PLA, in unpermeablized Calu-3 cells, Plscr1 and Ifn-λr1 colocalized on the cell membrane after IAV infection (Figure 4C). In contrast, neither colocalization nor clear expression of Plscr1 or Ifn-λr1 was evident without infection, implying an infection-specific interaction in human airway epithelial cells (Figure 4C). Hence, Plscr1 may regulate Ifn-λr1 expression and IFN-λ signaling in airway epithelial cells through both gene transcription and protein interaction in IAV infection.

**Figure 4.**
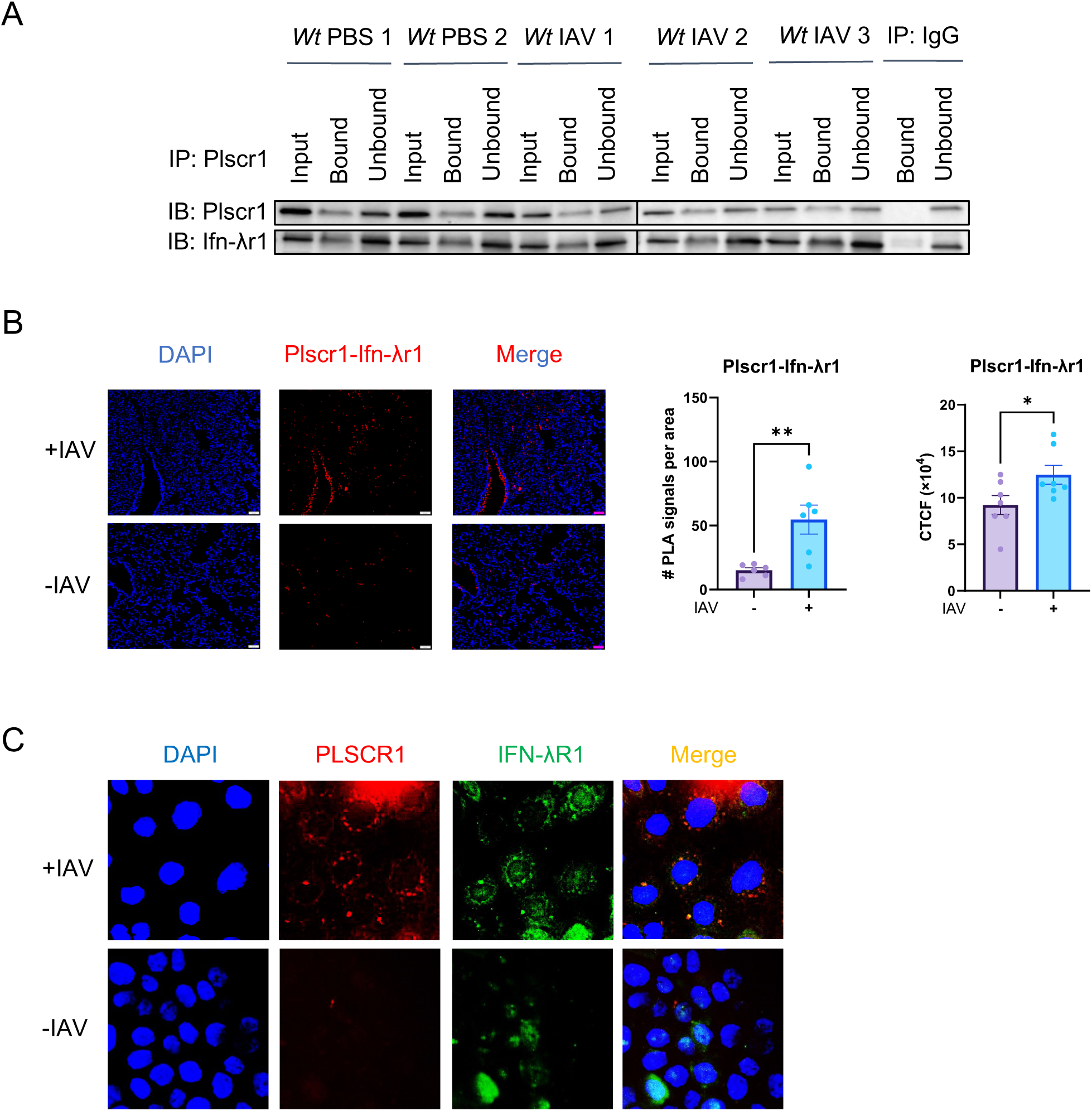
Protein Interaction between IFN-λR1 and PLSCR1 in IAV Infection. (A) Co-Immunoprecipitation of Plscr1 and Ifn-λr1 in whole mouse lungs followed by western blot. (B) Proximity ligation assay of Ifn-λr1 and Plscr1 in the lungs of *Wt* mice infected or uninfected with IAV. Scale bars represent 50 μm. Quantifications were performed using ImageJ. (C) Colocalization of IFN-λR1 (green) and PLSCR1 (red) on Calu-3 cell membranes infected or uninfected with IAV in a nonpermeabilized staining. Scale bars represent 10 μm. Data are expressed as mean ± SEM of n = 6-7 lungs/group. All data were pooled from three independent experiments. *p < 0.05, **p < 0.01. PLA, Proximity Ligation Assay. CTCF, Corrected Total Cell Fluorescence.

### Both Cell Surface and Nuclear PLSCR1 Regulates IFN-**λ** Signaling and Limits IAV Infection Independent of Its Enzymatic Activity

PLSCR1 mutants with specific PLSCR1 cellular distribution were employed to determine the relative contributions of cell surface and nuclear PLSCR1 in regulating IFN-λ signaling and IAV infection. PLSCR1 contains a 5-cysteine palmitoylation motif (C^184^CCPCC^189^), where substitution of these cysteines with alanine (labeled 5CA) completely abolishes its membrane localization, leading to exclusive localization in the cytosol and nucleus[26]. On the other hand, a single amino acid mutation of histidine^262^ to tyrosine (labeled H262Y) in the non-classical nuclear localization signal of PLSCR1 completely abates its nuclear localization, leaving PLSCR1 exclusively in the cytosol and on the cell membrane[27]. In addition, given that PS externalization plays a role in cell death regulation[19], we asked whether the enzymatic activity of PLSCR1 interferes with IFN-λ signaling. Phenylalanine^281^ in the β-barrel hydrophobic loop region of PLSCR1 has been shown to be important for Ca^2+^-dependent PS exposure[27], and a substitution with alanine (labeled F281A) renders PLSCR1 enzymatically inactive [15].

*PLSCR1(WT), PLSCR1(5CA), PLSCR1(H262Y)* and *PLSCR1(F281A)* plasmids built on PLV-EF1a-IRES-Hygro backbone were amplified in DH5α competent cells, co-transfected with HIV lentivector into 293T cells, and transduced into *Plscr1^-/-^* A549 cells. After a 10-day hygromycin selection, transduction efficiency and subcellular locations of PLSCR1 were analyzed using flow cytometry (Supplemental Figure 4A). *PLSCR1(WT)* exhibited higher PLSCR1 expression than *Plscr1^+/+^* cells, indicating overexpression resulting from transduction (Supplemental Figure 4B). Consistent with previous findings, *PLSCR1(5CA)* exhibited low surface expression, and *PLSCR1(H262Y)* had low nuclear expression of PLSCR1. *PLSCR1(F281A)* showed a similar distribution compared to *PLSCR1(WT)*, confirming that the loss of enzymatic activity did not restrict the subcellular localization. Transduction efficiency ranged between 37% to 70% (Supplemental Figure 4C).

In *PLSCR1^+/+^* cells or cells transduced with *PLSCR1(WT)*, *PLSCR1(5CA)* or *PLSCR1(F281A)*, IFN-λR1 expression was significantly induced by IAV infection. We found that PLSCR1 nuclear import was required for IFN-λR1 expression, as *IFN-*λ*R1* transcription and protein translation was not induced in cells transduced with *PLSCR1(H262Y)* or in *PLSCR1^-/-^* cells (Figure 5A and 5B). In contrast, minimal protein interactions between PLSCR1 and IFN-λR1 were detected by PLA in cells transduced with *PLSCR1(5CA) or in PLSCR1^-/-^ cells,* while *PLSCR1^+/+^* cells and cells transduced with *PLSCR1(WT)*, *PLSCR1(H262Y)* and *PLSCR1(F281A)* had strong protein binding of PLSCR1-IFN-λR1, demonstrating that PLSCR1 interacts with IFN-λR1 on cell membrane (Figure 5C). Furthermore, the lipid scramblase activity of PLSCR1 is uncoupled from its regulation of IFN-λ signaling.

**Figure 5.**
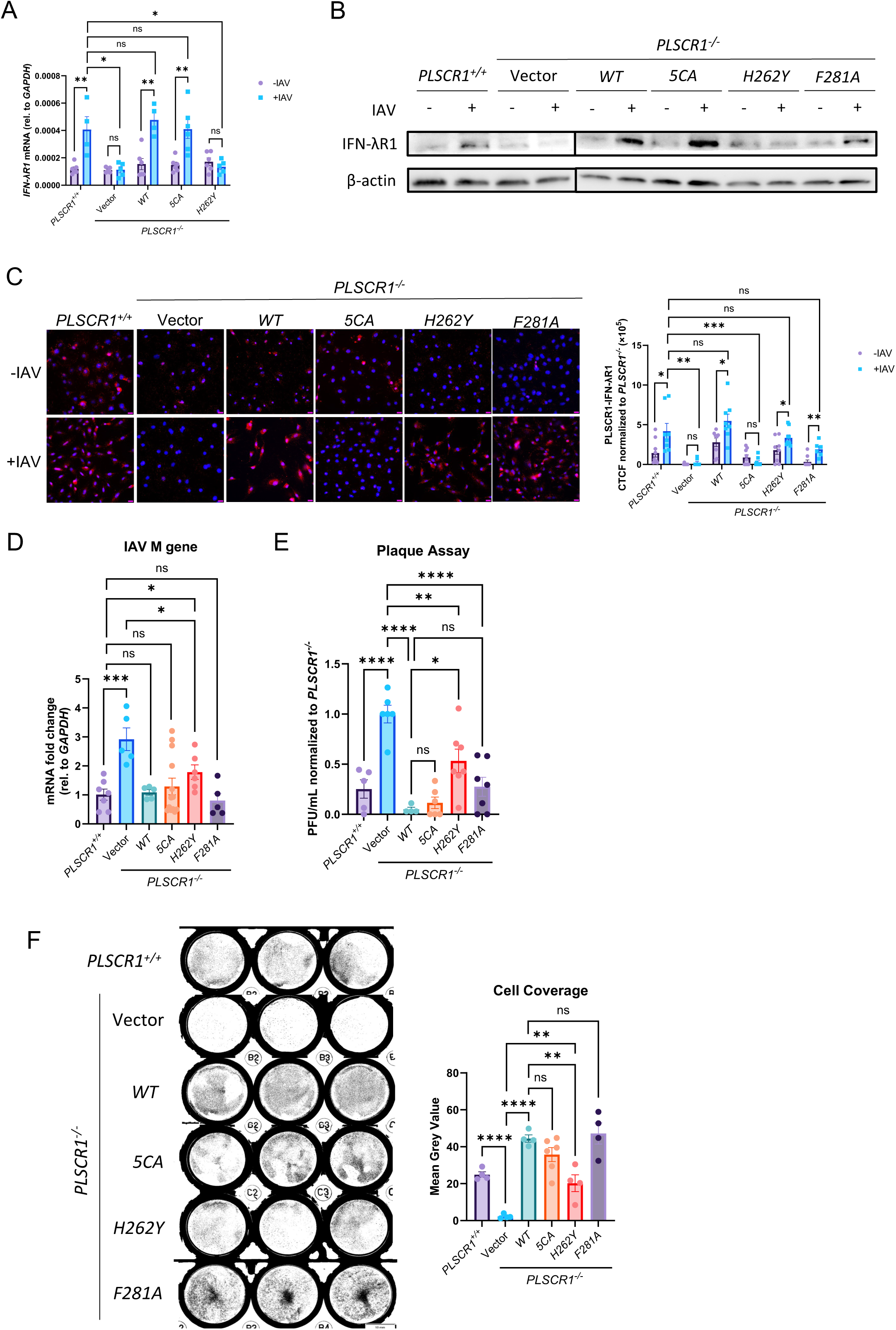
Requirement of Both Nuclear and Surface PLSCR1 but Not the Enzymatic Activity in IFN-λR1-Mediated Anti-Influenza Activities. *PLSCR1^-/-^* A549 cells were transduced with mutated *PLSCR1* plasmids using lentivirus and infected with IAV (PR8) for 24 hours at 1 MOI (A-E) or 10 MOI (F). (A) *IFN-*λ*R1* RNA by qRT-PCR. (B) IFN-λR1 proteins by western blotting. (C) Proximity ligation assay of IFN-λR1 and PLSCR1. Scale bars represent 20 μm. Quantifications were performed using ImageJ. (D) Viral RNA load was assessed by quantifying M gene by qRT-PCR. (E) Infectious viral titer was assessed by plaque assays. (F) Cells were stained with crystal violet. Cell viability was quantified using ImageJ. Data are expressed as mean ± SEM of n = 4-13 wells/group. All data were pooled from three independent experiments. ns, not significant, *p < 0.05, **p < 0.01, ***p<0.001, ****p<0.0001, *****p<0.00001. CTCF, Corrected Total Cell Fluorescence.

When compared to *PLSCR1^+/+^* cells, *PLSCR1(WT)*, *PLSCR1(5CA)* and *PLSCR1(F281A)* provided similar protection against IAV infection. However, *PLSCR1(H262Y)* showed only partial viral clearance, as demonstrated by viral copy numbers measured by M gene segment qRT-PCR quantification (Figure 5D) and infectious viral titers from plaque assays (Figure 5E). This is further validated with a cell coverage assay when a high dose IAV was used for infection. Since the proliferation between differently transduced cell lines was similar and all cells were grown to full confluency before infection, the cell coverage represented resistance to infection and survival of cells. While *PLSCR1(5CA)* and *PLSCR1(F281A)* had a high surface coverage comparable to *PLSCR1(WT)*, *PLSCR1(H262Y)* lost about half of the coverage (Figure 5F). Therefore, nuclear PLSCR1 is both essential and sufficient for IAV control through *IFN-*λ*R1* transcription regulation. Nevertheless, *PLSCR1(H262Y)* still rescued a significant number of cells when reintroduced into *PLSCR1^-/-^* cells, indicating a partial protective role of PLSCR1 on epithelial cell surface through IFN-λR1 protein regulation. Moreover, the anti-IAV function of PLSCR1 is independent of its lipid scramblase activity. Importantly, our findings that the scramblase activity is not required for protection against IAV infection is consistent with earlier observations with SARS-CoV-2[15], suggesting that the scramblase activity of PLSCR1 may generally be dispensable for protection against major human viruses.

### The Anti-Influenza Activity of Plscr1 Is Highly Dependent on Ifn-**λ**r1

To establish a causal link between the impaired type 3 IFN pathway and the increased susceptibility to IAV observed in *Plscr1^-/-^*mice and *PLSCR1^-/-^* cells, *Ifn-*λ*r1^-/-^* mice were crossed with *Plscr1^-/-^* mice to generate double-knockout *Plscr1^-/-^Ifn-*λ*r1^-/-^* mice. Immunofluorescence confirmed the absence of Ifn-λr1 expression in *Ifn-*λ*r1^-/-^* mice (Supplemental Figure 5A). Following sublethal IAV infection, *Ifn-*λ*r1^-/-^* mice exhibited significantly greater body weight loss at 3 dpi than *Plscr1^-/-^*mice, which retain basal Ifn-λr1 expression (Supplemental Figure 5B). Moreover, they also showed significantly higher total BAL cell counts (Supplemental Figure 5C), increased neutrophil percentages (Supplemental Figure 5D), and higher infectious viral titers as measured by plaque assays (Supplemental Figure 5E). These findings indicate that complete loss of Ifn-λr1 results in greater susceptibility to IAV than the loss of Plscr1-mediated Ifn-λr1 upregulation, underscoring the essential role of the type 3 IFN pathway in anti-viral defense.

Importantly, *Plscr1^-/-^Ifn-*λ*r1^-/-^* mice showed no further increase in weight loss (Supplemental Figure 5B), total BAL cell counts (Supplemental Figure 5C), neutrophil percentages (Supplemental Figure 5D), and IAV titers (Supplemental Figure 5E) compared to *Ifn-*λ*r1^-/-^* mouse lungs, indicating that the antiviral activity of Plscr1 is largely dependent on Ifn-λr1.

### PLSCR1 Expression Is Upregulated in the Ciliated Airway Epithelial Compartment of Mice following Flu Infection

To understand the specific cell types that have increased PLSCR1 expression following flu infection *in vivo*, we explored a comprehensive scRNA sequencing dataset generated from uninfected or IAV-infected mouse lungs at 0, 1, 3, 6 and 21 dpi[28]. Distinct lung cell populations were identified using the Seurat algorithm. We assigned a total of 38 clusters based on their transcriptomic signatures, including 13 epithelial populations, 5 endothelial populations, 9 mesenchymal populations and 11 immune populations (Table S2). Two-dimensional UMAP (Figure 6A) and bar charts of the proportion (Supplemental Figure 6A) and cell count (Supplemental Figure 6B) of each cluster demonstrated that these main lung cell populations (epithelial, endothelial, mesenchymal, and immune) were dynamic over the course of infection. Following infection, many populations emerged, particularly within the immune cell clusters. At the same time, some clusters were initially depleted and later restored, such as microvascular endothelial cells (cluster 2). Other populations, such as interferon-responsive fibroblasts (cluster 20), showed a dramatic yet transient expansion during acute infection and disappeared after infection resolved.

**Figure 6.**
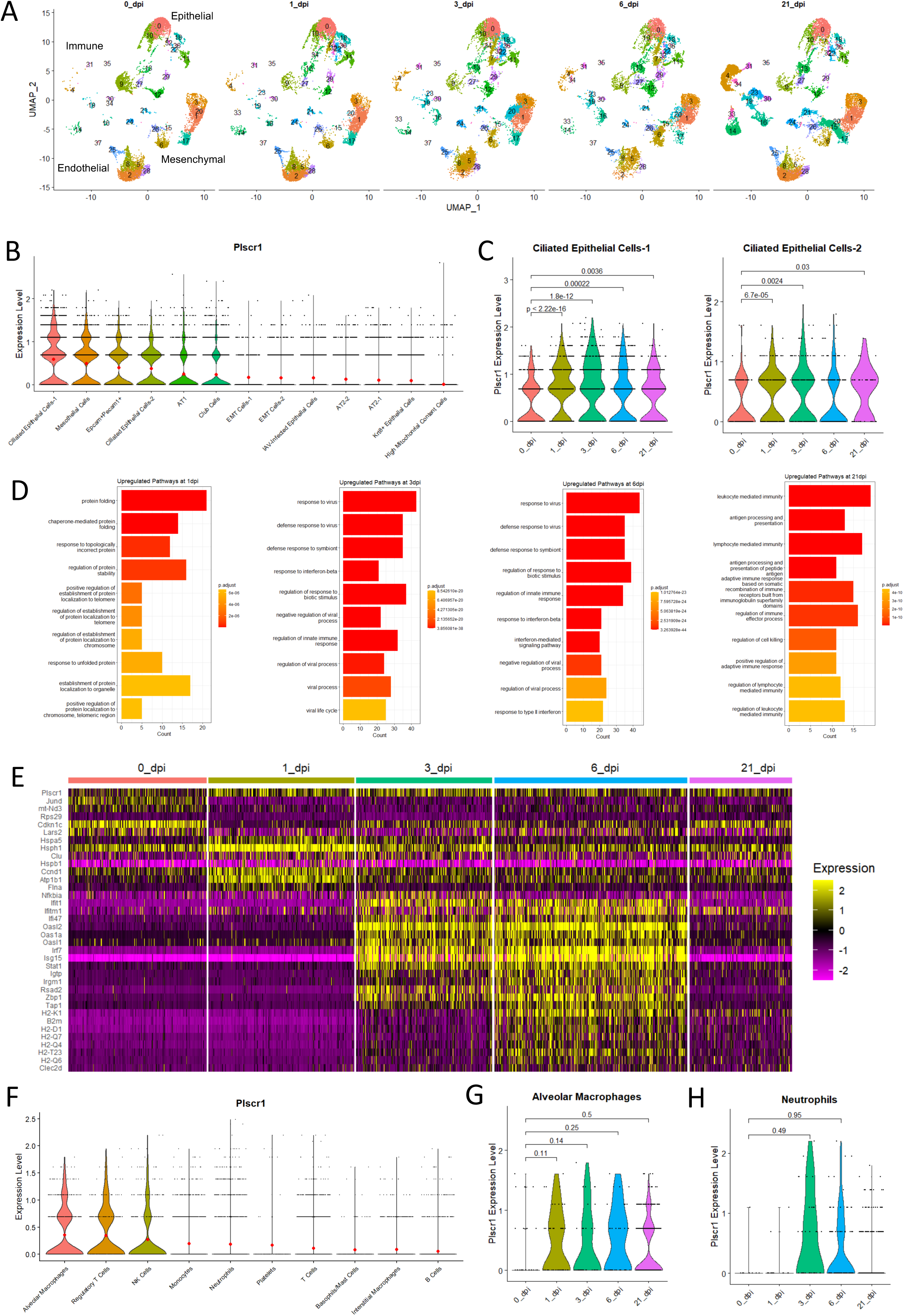
Cell-Specific Roles of *Plscr1* in Influenza Virus Infection in Mice. *Wt* mice were exposed to 2500 EID50 IAV (PR8) infection. Lungs were used for single-cell RNA sequencing analysis at 0, 1, 3, 6 and 21 dpi. (A) Two-dimensional UMAP representation of individual cells obtained from different timepoints. (B) Violin plot of aggregated *Plscr1* expressions in all epithelial cell clusters. Red dots represent mean expression levels. (C) Violin plot of time-dependent *Plscr1* expressions in both ciliated epithelial cell clusters. (D) GO analysis for upregulated pathways in Ciliated Epithelial Cells-1. (E) Heatmap of the most differentially expressed genes of Ciliated Epithelial Cells-1 at different timepoints. (F) Violin plot of aggregated *Plscr1* expressions in all immune cell clusters. Red dots represent mean expression levels. (G) Violin plot of time-dependent *Plscr1* expressions in alveolar macrophage cluster. (H) Violin plot of time-dependent *Plscr1* expressions in neutrophil cluster.

Within the epithelial populations, *Plscr1* was mainly expressed by ciliated epithelial cells, mesothelial epithelial cells, Epcam+Pecam1+ cells, club cells and AT1 cells (in decreasing order of aggregated expression, Figure 6B). Conversely, AT2 cells, epithelial-mesenchymal transitional cells and Krt8+ cells exhibited very low levels of *Plscr1* expression. Among the various epithelial populations, ciliated epithelial cells exhibited both the highest aggregated expression of *Plscr1* (Figure 6B) and the most significant upregulation (p < 2.22e-16 and p = 6.7e-05) at 3 dpi in early IAV infection (Figure 6C and Supplemental Figure 7A-7K). In contrast, AT1 cells were the only other epithelial cluster to show *Plscr1* upregulation at 3dpi, but to a much less extent (p = 0.033, Supplemental Figure 7J). Moreover, the increased expression of Plscr1 persisted for at least 21 days in ciliated epithelial cells, suggesting that Plscr1 may primarily exert its anti-flu activities in these cells (Figure 6C).

**Figure 7.**
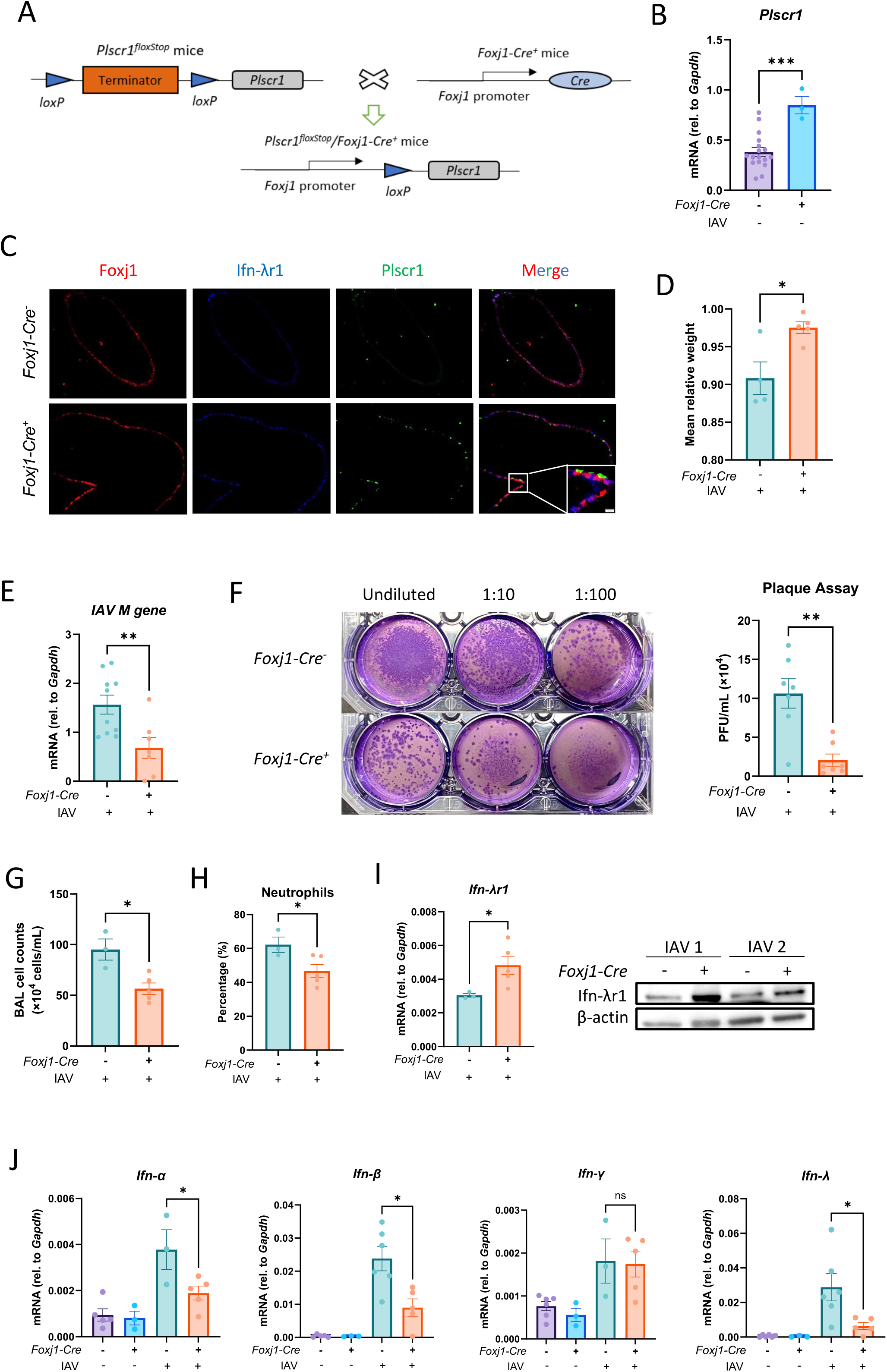
Reduced Susceptibility of *Plscr1^floxStop^Foxj1-Cre^+^* Mice to Influenza Virus Infection. *Plscr1^floxStop^* and *Plscr1^floxStop^Foxj1-Cre^+^* mice were exposed to sublethal (300 pfu) IAV (WSN) infection and sacrificed at 3 dpi. (A) Schematic representation of the experimental design of of ciliated epithelial cell conditional *Plscr1 KI* mice. (B) Validation of *Plscr1* overexpression in lungs of *Plscr1^floxStop^Foxj1-Cre^+^* mice by qRT-PCR. (C) Representative immunofluorescent staining for Plscr1, Ifn-λr1 and Foxj1 in lungs. (D) Mean relative weight of mice. (E) Viral RNA load in the lungs was assessed by quantifying M gene by qRT-PCR. (F) Infectious viral titer in the lungs was assessed by plaque assays. (G) Total BAL leukocyte numbers. (H) Neutrophil percentages in BAL. (I) Whole lungs were analyzed for *Ifn-*λ*r1* RNA by qRT-PCR and Ifn-λr1 protein by western blot. (J) Whole lungs were analyzed for *Ifn-*α, *Ifn-*β, *Ifn-*γ, *Ifn-*λ RNA by qRT-PCR. (K) Model depicting proposed mechanism of PLSCR1-regulated IFN-λ signaling. Data are expressed as mean ± SEM of n= 3-10 mice/group. ns, not significant, *p < 0.05, **p < 0.01, ***p<0.001. dpi, days post infection.

For the predominant ciliated epithelial cell cluster, we performed Gene Ontology (GO) analysis of the differentially expressed genes to compare potentially distinct functions before and after infection (Figure 6D). We found that the most upregulated pathways at 1 dpi were related to protein folding and localization, indicating a rapid cellular response to infection by altering their protein profile. At 21 dpi, ciliated epithelial cells were characterized by adaptive immune regulation, such as antigen presentation and memory cell generation. At 3 and 6 dpi, ciliated epithelial cells exhibited similar innate immune and inflammatory signatures dominated by interferon signaling. As expected, *Plscr1* participated in all of the top 10 enriched pathways, implying that it could facilitate robust antiviral responses in ciliated epithelial cells as an ISG during acute infection.

Gene expression levels of the most differentially expressed genes at individual timepoints in ciliated epithelial cells were shown in a heatmap (Figure 6E). Signature genes for cells at homeostasis included *Jund* (JunD proto-oncogene), *mt-Nd3* (mitochondrial NADH dehydrogenase 3) and *Cdkn1c* (cyclin-dependent kinase inhibitor 1C), which are regulators of cell metabolism and proliferation[29–31]. Signature genes enriched at 1 dpi were *Hspa5* (heat shock protein family A member 5), *Hsph1* (heat shock protein family H member 1) and *Clu* (clusterin), which play critical roles in protein folding and assembly[32–34]. Consistent with the GO analysis, ciliated cells at 3 and 6 dpi had very similar signature genes associated with interferon pathways, namely a large component of ISGs, such as *Isg15* (interferon-stimulated gene 15), *Oasl2* (2’-5’-oligoadenylate synthetase like 2) and *Irf7* (interferon regulatory factor 7). Importantly, Plscr1 is required to potentiate the expressions of many of these ISGs[23], highlighting its potentially multifunction in anti-flu responses of ciliated epithelial cells. Signature genes for 21 dpi largely overlapped with those for 10 dpi, containing many histocompatibility 2, class II antigen proteins, which are mediators of MHC class II antigen processing and presentation.

In contrast to epithelial clusters, immune populations had low levels of *PLSCR1*, with relatively high expressions observed in alveolar macrophages, NK cells and regulatory T cells (Figure 6F). However, we did not observe any significant difference in PLSCR1 levels before and post-infection in macrophages and neutrophils (Figure 6G and 6H), the two immune populations that express IFN-λR1 besides dendritic cells[35]. Hence, the anti-influenza activities we described are unlikely to be carried out by immune cells.

### Overexpression of Plscr1 in Ciliated Epithelial Cell, but Not Myeloid Cell, Is Sufficient to Provide Anti-Flu Protection through IFN-**λ** Signaling

To further validate the relevant contributions of ciliated epithelial cells versus myeloid populations including macrophages and neutrophils, we generated *Rosa26* locus targeted *Plscr1* conditional knock-in transgenic mice (*Rosa26-LoxP-STOP-LoxP-Plscr1 Tg*; labeled *Plscr1^floxStop^*). These mice were bred with *Foxj1-Cre* or *LysM-Cre* mice to overexpress Plscr1 in ciliated epithelial cells or myeloid cells respectively (Figure 7A). Overexpression was confirmed by qRT-PCR with whole lung lysate and immunofluorescence staining at baseline (Figure 7B, 7C and Supplemental Figure 8A). At 3 dpi with sublethal IAV infection, *Plscr1^floxStop^Foxj1-Cre^+^*mice lost less body weight (Figure 7D). Moreover, they had lower viral copy numbers as measured by IAV M gene segment qRT-PCR (Figure 7E), and lower infectious viral titer measured by plaque assays (Figure 7F) compared to *Plscr1^floxStop^* mice, indicating overexpression of Plscr1 in ciliated epithelial cells provides protection against flu infection. Concurrently, they had significantly lower total BAL cell counts (Figure 7G), lower neutrophil percentages (Figure 7H) and lower type 1 and 3 *Ifn* expressions (Figure 7J). We then examined Ifn-λr1, and found that its transcription and protein expression were further increased in *Plscr1^floxStop^Foxj1-Cre^+^* mice during acute infection (Figure 7I). Therefore, ciliated epithelial cell-specific overexpression of Plscr1 is sufficient to provide anti-influenza protection through IFN-λ signaling. In contrast, using similar approaches, we found that overexpression of Plscr1 in myeloid cells did not protect mice from IAV infection: *Plscr1^floxStop^LysM-Cre^+^*mice exhibited comparable weight loss, total BAL cell counts, viral copy numbers, histopathology and expressions of interferons, *Ifn-*λ*r1* and *Plscr1* at both early and late timepoints examined (Supplemental Figure 8B-8I).

**Figure 8.**
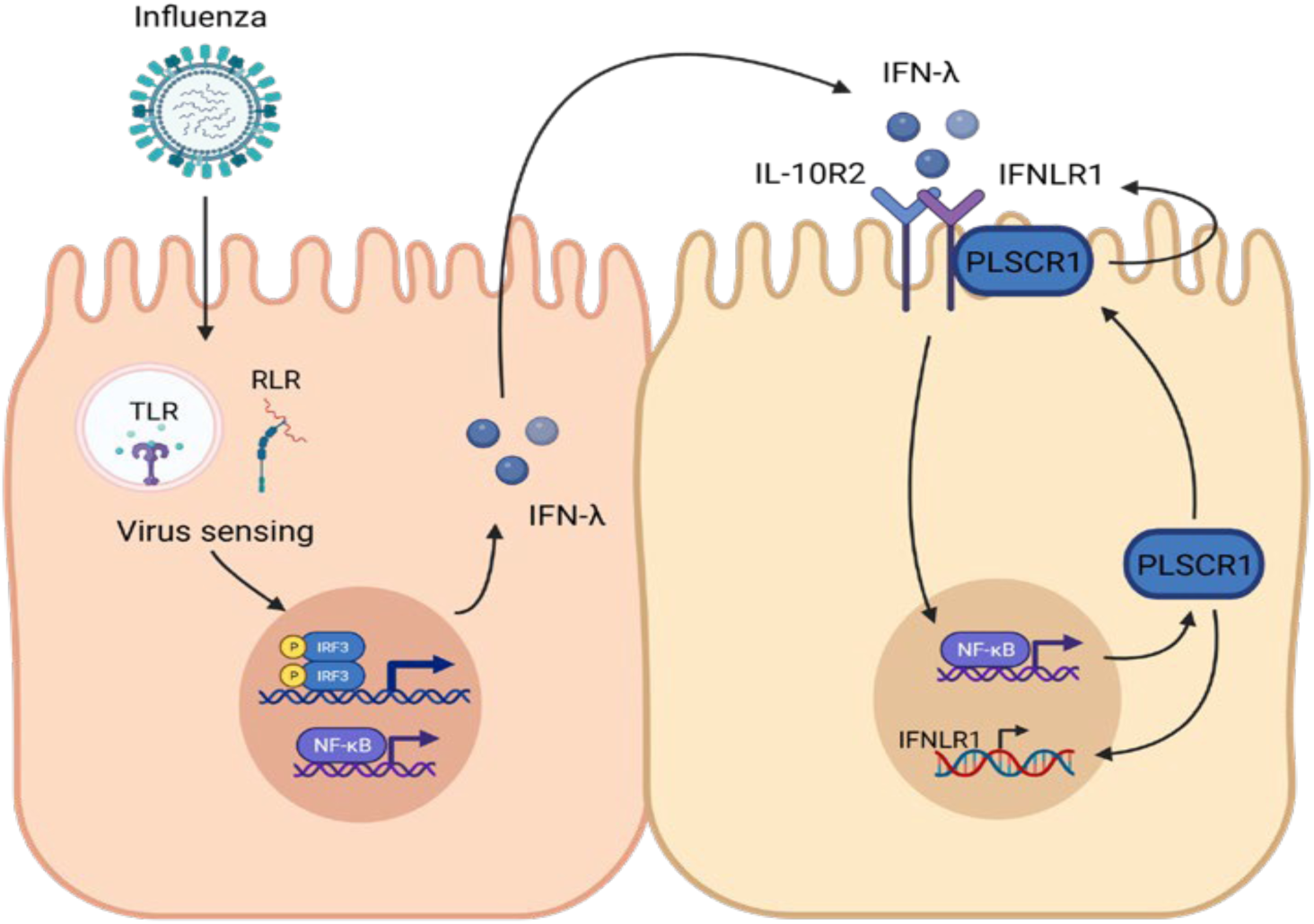
Model depicting proposed mechanism of PLSCR1-regulated IFN-λ signaling. Influenza infection is usually detected by intracellular pattern recognition receptors (PRRs) such TLR 3 and 7, RIG-I and MDA5. These PRRs activate the expression of *IFN-λ* in early infection stage through IRF-3 and NK-κB-controlled transcriptions. IFN-λ secreted by the infected cells interacts with IL-10R2 and IFN-λR1 on neighboring cell surface, which results in activation of expression of various IFN-stimulated genes, including *PLSCR1*. In ciliated airway epithelial cells, PLSCR1 can further enhance the transcription of *IFN-λR1* by directly binding to its promoter region as a transcriptional factor, or interact with IFN-λR1 on the cell membrane.

Taken together, consistent with the scRNA sequencing, transgenic mouse models with cell-specific overexpression of Plscr1 demonstrated that Plscr1 offers antiviral protection in ciliated airway epithelial cells through IFN-λ signaling (Figure 8).

## DISCUSSION

We are the first group to demonstrate the roles of Plscr1 in a mouse-adapted human IAV-infected mouse model, to implicate its IFN-λ signaling-related mechanisms, and to elucidate the cell types that are responsible for Plscr1-mediated anti-influenza activities. We established *Plscr1^-/-^* mice and found them more susceptible to IAV (WSN) compared to *WT* mice, as evidenced by greater weight loss in both sublethal and lethal infection and poorer survival in a lethal infection. Further examination of infected lungs provided the first *in vivo* evidence demonstrating that Plscr1 suppressed human IAV replication. This observation aligns with previous report indicating that PLSCR1 interacts with the IAV NP, thereby impairing its nuclear import *in vitro*[20]. Notably, while differences in viral copy numbers were only observed at the early stages of infection, coinciding with a significant increase in *Plscr1* transcription, these changes had profound implications for host fitness. Therefore, as one of the earliest induced ISGs, Plscr1 constitutes the frontline defense against influenza infection.

While the only previously published *Plscr1^-/-^* mouse flu model focused on an H1N1 SIV infection[21], our data showed both similarities and discrepancies. First, while both studies observed that Plscr1 promoted survival during IAV infection, SIV-infected *Plscr1^-/-^*mice exhibited weight loss similar to *Wt* mice. Furthermore, while both models attributed the lower survival rate in *Plscr1^-/-^* mice to increased viral replication, SIV-infected *Plscr1^-/-^* lungs exhibited higher viral titers across all examined time points, from 1 to 7 dpi. Intriguingly, contrary to our observations, *Plscr1* expression was markedly decreased in SIV infection. Given previous *in vitro* studies demonstrating PLSCR1 induction by IAV (WSN)[20] and type 1 IFNs[13, 23, 36], we propose that the contradictory trend observed by Liu et al. may be attributed to distinct properties of SIV, such as viral replication rate, both the cellular tropism and the tissue tropism (proximal or distal lung), or antigen variation which may affect direct interaction with PLSCR1, innate sensing of the infection, or recognition by the adaptive immune response.

The delicate balance between immunity and immunopathology plays a pivotal role in determining host fitness during viral infections. To interrogate immunopathology in the lungs, we accessed the BAL, histology and interferon expressions. BAL from *Plscr1^-/-^* mice were highly enriched with inflammatory neutrophils and lymphocytes, which were likely attracted by robust IFNs and other chemokines. Consistently, *Plscr1^-/-^*mice exhibited more severe lung damage and a greater extent of affected areas. These findings indicate that Plscr1 not only enhances immunity but also mitigates immunopathology. Importantly, regardless of excessive production of antiviral IFNs in *Plscr1^-/-^* mice, they failed to effectively control the initial viral infection. This suggests that the absence of Plscr1 impairs the IFN signaling pathway, highlighting the crucial role of Plscr1 in facilitating effective antiviral responses.

Although type 1 and 3 IFNs may share similar downstream pathways, they rely on distinct receptors for signaling. Consistent with previous findings[3], Ifn-λr1 was detected in respiratory epithelium, including ciliated epithelial cells, club cells and AT2 cells during infection. Loss of Plscr1 impaired *Ifn-*λ*r1* transcription in IAV infection, with this transcriptional difference translating into protein expression. IFN-λ is crucial for early viral control within the initial days of infection without igniting unnecessary inflammation and compromising host fitness[10]. With limited Ifn-λr1 expression, *Plscr1^-/-^* mice were unable to mount a robust type 3 IFN response to control early viral infection. Instead, they relied largely on type 1 interferons, which succeeded in eliminating IAV at later time points, but led to exaggerated immunopathology. Furthermore, our observations of enhanced neutrophilia, lung injury and lethality in *Plscr1^-/-^*mice align with findings reported in *Ifn-*λ*r1^-/-^*mice in IAV infection [10]. However, a discrepancy in *Ifn-*λ*r1* expression over the course of infection was observed between the RNA sequencing and the qRT-PCR data. While RNA-seq showed further upregulation of *Ifn-* λ*r1* at 7 dpi (Figure 3A), qRT-PCR indicated a rapid downregulation at the same time point (Figure 3B). The reason for this time-dependent discrepancy remains unclear and warrants further investigation. In addition, this study did not definitively establish causality between reduced Ifn-λ signaling and the observed *in vivo* phenotype. The increased morbidity and mortality in *Plscr1^-/-^* mice could also be attributed to elevated Tnf-α levels and associated lung damage. Given that proinflammatory cytokines and/or enhanced lung damage are known contributors to influenza morbidity and mortality, future work will be needed to disentangle the impacts of TNF-α, IL-1β and other inflammatory cytokines from those of the IFN pathway to fully clarify the role of Plscr1 in antiviral defense.

PLSCR1 expression was increased in response to IFN-λ in human airway epithelial cells, consistent with previous studies[15]. While PLSCR1 typically localizes on the cell membrane and in the cytoplasm, it translocates into nucleus to bind the *IFN-*λ*R1* promoter upon IAV infection, thereby regulating *IFN-*λ*R1* transcription. The nuclear localization and functions of PLSCR1 have been extensively documented in previous studies[17, 37–39]. Relevantly, IFN-α promotes the nuclear translocation of PLSCR1 in breast cancer cells[26]. Therefore, it is highly plausible that the nuclear trafficking of PLSCR1 in airway epithelial cells is similarly stimulated by IFN-α produced during IAV infection, but further evidence is demanded. Additionally, the precise binding site for PLSCR1 within the *IFN-*λ*R1* promoter and the binding motif on PLSCR1 remain unknown. Previous bioinformatics predictions revealed that the −430∼-421 segment of *IFN-*λ*R1* promoter likely contains binding sites for a number of transcription factors (TFs) with important regulatory functions[40]. Further mutagenesis studies, such as truncations or single nucleotide mutations within these sequences, could be done to identify the specific motif for PLSCR1 binding. Finally, it is not clear whether PLSCR1 directly activates *IFN-*λ*R1* expression by acting as a TF, or as a co-factor enhancing other TF’s transcriptional activity. Coimmunoprecipitations could be pursued in future to explore the binding potential between PLSCR1 and other known TFs for *IFN-*λ*R1*, such as NF-Y[40].

In addition to activities in the nucleus, PLSCR1 has been shown to interact with multiple proteins on the plasma or endosomal membrane[22, 41–44]. Here we reported a novel interaction between PLSCR1 and IFN-λR1 on airway epithelial cell membrane *in vivo* and *in vitro*, confirmed with both coimmunoprecipitation and immunofluorescence. Since their interaction was significantly enhanced after IAV infection, we speculate that membrane-bound PLSCR1 is a positive regulator of IFN-λR1 signaling. One plausible mechanism is that PLSCR1 facilitates the intracellular trafficking of IFN-λR1, akin to its role in assisting the trafficking of other membrane receptors[22, 42, 45].

The subcellular location of PLSCR1 is vital not only for interactions with host components, but also for direct viral control. We found that nuclear PLSCR1 is both necessary and sufficient for viral control in airway epithelial cells, whereas membrane PLSCR1 provides only partial protection against IAV infection. These findings are not surprising, as the previously reported anti-flu mechanism of PLSCR1 also relies on its nuclear localization signal to restrict the import of IAV NP[20]. Furthermore, besides *IFN-*λ*R1*, PLSCR1 enhances the expression of a select subset of ISGs in IAV infection as well, a mechanism potentially mediated by nuclear PLSCR1 on ISG gene transcription[23]. On the other hand, membrane PLSCR1 may modulate the JAK/STAT signaling pathway, thereby augmenting the optimal anti-viral activity of these ISGs[23]. We found that membrane PLSCR1 interacts with IFN-λR1 protein in IAV infection, suggesting that it could facilitate viral elimination to some extent.

PLSCR1 is most well-known for its scramblase activity that favors PS exposure, apoptosis and phagocytosis[41, 46]. Using an enzymatically inactive mutant of PLSCR1, we uncoupled its lipid scramblase activity from anti-influenza activity. There are several potential explanations for this finding. First, our epithelial cell culture lacked phagocytes, therefore the impact of apoptosis followed by phagocytosis induced by PLSCR1 is minimal. Future studies using mice that harbor *Plscr1(F281A)* mutation would be needed to verify the role of lipid scramblase activity and epithelial cell apoptosis in the presence of phagocytes. Second, PLSCR1 exhibits only weak enzymatic activities compared to other members of lipid scramblase family, possibly due to its vastly different central β-barrel structure[15, 47]. PS externalization may be compensated by other more potent scramblases. Importantly, the lipid scramblase activity of PLSCR1 has been shown to be dispensable for its anti-SARS-CoV-2 function in a similar manner[15], suggesting a general lack of significance for its enzymatic activity in viral infections.

Although PLSCR1 has several previously described anti-influenza functions, including interfering with viral nuclear import[20], regulating TLR9 signaling [22], and potentiating the expression of other ISGs[23], our studies have clarified the relative contribution of the type 3 IFN pathway to Plscr1-mediated anti-influenza immunity using *Plscr1^-/-^Ifn-*λ*r1^-/-^*mice. We observed that *Ifn-*λ*r1^-/-^* mice were more susceptible to IAV infection than *Plscr1^-/-^*, suggesting that the complete loss of Ifn-λr1 results in worse protection than impaired Ifn-λr1 upregulation alone. Moreover, the previously identified anti-IAV functions of Plscr1 do not appear sufficient to compensate for the loss of Ifn-λr1 signaling in *Ifn-*λ*r1^-/-^*mice. The absence of further disease exacerbation or increased viral titers in *Plscr1^-/-^Ifn-*λ*r1^-/-^* mice compared to *Ifn-*λ*r1^-/-^* mice indicates that the anti-influenza activity of Plscr1 is largely dependent on Ifn-λr1.

While scRNA-seq analysis revealed that endothelial cells express Plscr1 most abundantly in the lung, they are not the major target of IAV infection, and IAV does not efficiently replicate in them[48]. Instead, airway epithelial cells are the frontline defense against respiratory pathogens, with ciliated epithelial cells being the only cell type that express α2,3-linked SA, the primary influenza virus receptor in the mouse airway[49]. Coincidently, our scRNA-seq results showed that ciliated epithelial cells not only had the highest aggregated expression of *Plscr1*, but also had the most significant increase in *Plscr1* expression in early IAV infection at 3 dpi. Experiments with *Plscr1^floxStop^Foxj1-Cre^+^*mice further supported ciliated epithelial cell-dependent protection against IAV, with improved immunity and viral clearance, and dampened immunopathology at as early as 3 dpi. These findings suggest that as a result of enhanced Ifn-λr1 due to Plscr1 overexpression, type 3 interferons were able to exert their advantages being the earliest produced interferon, mounting both antiviral and anti-inflammatory responses in ciliated epithelial cells. To further establish the causal relationship between Plscr1 and Ifn-λ signaling in airway ciliated epithelial cells, future experiments should focus on specifically overexpressing Plscr1 in ciliated epithelial cells on an *Ifn-*λ*r1^-/-^* background by breeding *Plscr1^floxStop^Foxj1-Cre^+^Ifn-*λ*r1^-/-^* mice. In addition, ciliated epithelial cells isolated from *Ifn-*λ*r1^-/-^*murine airways could be transduced with a *Plscr1* overexpression construct. We hypothesize that overexpression of Plscr1 in ciliated epithelial cells would not be able to rescue susceptibility in *Ifn-*λ*r1^-/-^* mice or cells, as the *Plscr1^-/-^Ifn-*λ*r1^-/-^* mouse model suggests that Plscr1’s Ifn-λr1-independent anti-influenza mechanisms are likely minor compared to its role in upregulating Ifn-λr1.

Taken together, our findings highlight the essential role of PLSCR1 in the regulation of IFN-λR1 transcription in nucleus and expression on plasma membrane, both *in vitro* and *in vivo*. These mechanisms are crucial for inhibiting viral spread, reducing inflammation, and enhancing overall host fitness during IAV infection. Further, we found that the enzymatic activity of PLSCR1 is dispensable for its anti-influenza function. Finally, ciliated airway epithelial cells are the primary cell type in the lung for mounting PLSCR1-mediated anti-influenza responses. The potential of PLSCR1 agonists that target ciliated airway epithelial cells as therapeutic treatments for influenza holds promise for future medical interventions. Moreover, our results have the potential to impact the classical yet evolving field of IFN signaling. Not only do these findings elucidate and expand our understanding of newly discovered IFN-λ signaling, but they also shed light on the specific cell types and conditions under which IFN-λ signaling is modulated. Given the significance of IFN-λ signaling in various infectious diseases, these insights may pave the way for innovative therapeutics approaches targeting corresponding regulatory molecules in the treatment of other microbial infections in addition to influenza.

## MATERIALS AND METHODS

### Viruses

Purified A/WSN/1933 and A/PR/8/1934 (H1N1) virus (IAV) was kindly provided by Dr. Amanda Jamieson, Brown University. MDCK cells were used for the preparation of virus stocks and for virus titration by plaque assay.

### Cell culture

Calu-3 cells were kindly provided by Dr. Suchitra Kamle, Brown University. Cells were maintained in Eagle’s Minimum Essential Medium (ATCC) with 10% fetal bovine serum (Gibco) and 1% penicillin-streptomycin (Sigma-Aldrich).

MDCK cells were kindly provided by Dr. Amanda Jamieson, Brown University. Cells were maintained in Dulbecco’s Modified Eagle Medium (Genesee) with 10% fetal bovine serum and 1% penicillin-streptomycin.

*PLSCR1^-/-^* A549 cells were kindly provided by Dr. John MacMicking, Yale University. 293T cells were kindly provided by Dr. Suchitra Kamle, Brown University. Both cells were maintained in Dulbecco’s Modified Eagle Medium with 10% fetal bovine serum, 1% MEM non-essential amino acids (Gibco) and 1mM sodium pyruvate (Gibco).

### Mice

Wild-type (*Wt*) mice on a C57BL/6J genetic background were purchased from the Jackson Laboratories. *Plscr1^tm1a(EUCOMM)Hmgu^* mice (*Plscr1^-/-^* mice on C57BL/6 background) were purchased from the International Mouse Phenotyping Consortium. Those mice have an FRT flanked *lacZ*/neomycin sequence-tagged *LoxP* insertion upstream of *Plscr1* exon 6 and 7, and another *LoxP* insertion downstream of these critical exons. Subsequent *Cre* expression results in whole-body knockout mice that transcribed a shortened and nonfunctional transcript of *Plscr1*[50]. Genetic screening of *Plscr1^-/-^* mice was carried out by conventional PCR on genomic DNA from mouse tail biopsies using the following primers for *LacZ* reporter: Fw: 5’-GCGATCGTAATCACCCGAGT-3’ and Rev: 5’-CCGCCAAGACTGTTACCCAT-3’. This set of primers generates a 307bp fragment in *Plscr1^-/-^* mice.

*Rosa26* locus targeted *Plscr1* conditional knock-in transgenic mice (*Rosa26-LoxP-STOP-LoxP-Plscr1 Tg*; *Plscr1^floxStop^*mice on C57BL/6 background) were generated at Brown Mouse Transgenic and Gene Targeting Facility. They were bred with *LysM-Cre* mice purchased from the Jackson Laboratories and *Foxj1-Cre* mice gifted by Drs. Yong Zhang and Michael Holtzman at Washington University School of Medicine in St. Louis to generate *Plscr1^floxStop^LysM-Cre^+^* and *Plscr1^floxStop^Foxj1-Cre^+^*mice respectively. For the detection of *Cre* recombinase in *LysM-Cre* mice, the following primers were used: Common Fw: 5’-CTTGGGCTGCCAGAATTTCTC-3’, Mutant Rev: 5’-CCCAGAAATGCCAGATTACG-3’ and Wild type Rev: 5’-TTACAGTCGGCCAGGCTGAC-3’. A 700bp fragment is amplified in homozygous mutants, while a 350bp fragment is amplified in wild type mice. Both fragments show up in heterozygotes. For the detection of *Cre* recombinase in *Foxj1-Cre* mice, the following primers were used: Mutant Fw: 5’-CGTATAGCCGAAATTGCCAGG-3’ and Mutant Rev: 5’-CTGACCAGAGTCATCCTTAGC-3’. A 327bp fragment is amplified in homozygous mutants.

*Ifn-*λ*r1^tm1a(EUCOMM)Wtsi^*; Deleter-Cre mice (*Ifn-*λ*r1^-/-^*mice on C57BL/6 background) were generated and gifted by Dr. Sanghyun Lee at Brown University[51, 52]. They were bred with *Plscr1^-/-^*mice to generate *Plscr1^-/-^Ifn-*λ*r1^-/-^* mice. For the detection of *Ifn-*λ*r1* in those mice, the following primers were used: Common Fw: 5’-AGGGAAGCCAAGGGGATGGC-3’; Rev 1: 5’-AGTGCCTGCTGAGGACCAGGA-3’; Rev 2: 5’-GGCTCTGGACCTACGCGCTG-3’. A 564bp fragment is amplified in homozygous *Ifn-*λ*r1^-/-^* mutants, while a 231bp fragment is amplified in wild type mice.

All mice were housed and further bred in Brown University animal facilities. The mice with null mutation or overexpression of Plscr1 did not show any apparent abnormal phenotypes and developmental and signaling issues. All murine procedures were approved by the Institutional Animal Care and Use Committees at Brown University.

### Infection and treatment of mice

10-12-week-old mice were intranasally infected once with various doses of IAV in 30 μL of sterile PBS (Gibco), or treated with 2.5 μg/g of body weight of poly(I:C) (Invivogen) every day for 6 days under isoflurane-based anesthesia. Poly(I:C) was warmed up in a water bath at 37 °C before administration. Thirty μL of sterile PBS was administrated in control mice. Only males were used in infections. Littermates were randomly assigned to experimental and control groups.

### BAL harvest and differential cell counts

Mice were euthanized with intra-peritoneal injection of 300 μL of urethane (0.18 g/ml, Sigma-Aldrich). Bronchoalveolar lavage (BAL) of the whole lung was performed with a total of 1 mL ice-cold PBS via an incision at the trachea. An aliquot was stained with trypan blue solution (Gibco) for viability and cell number determination using TC20 automated cell counter (Bio-Rad). Samples were centrifuged at 2500 rpm for 5 min at 4°C. Cells were pelleted onto glass slides via cytospin centrifugation at 700 rpm for 7 min and stained with Hema 3 (Fisher Scientific). Neutrophils, macrophages, lymphocytes and eosinophils were counted under a light microscope.

### RNA isolation and qPCR

Harvested right lungs were immediately snap-frozen in liquid nitrogen and stored at - 80°C afterwards. Frozen lungs were homogenized in TRIzol Reagent (Invitrogen). Cells were washed with ice cold PBS once, directly lysed in the culture dish with TRIzol Reagent and collected using a cell scraper. RNA isolation was performed with RNeasy Mini Kit (Qiagen). 0.5 μg of the isolated RNA was used for cDNA synthesis with iScript cDNA Synthesis Kit (Bio-Rad). Real-time quantitative PCR was performed with iTaq Universal SYBR^®^ Green Supermix (Bio-Rad). All primers used were listed in Table S1. Relative amounts of mRNA expression were normalized to the level of mouse *Gapdh* or human *GAPDH*.

**Table S1.**
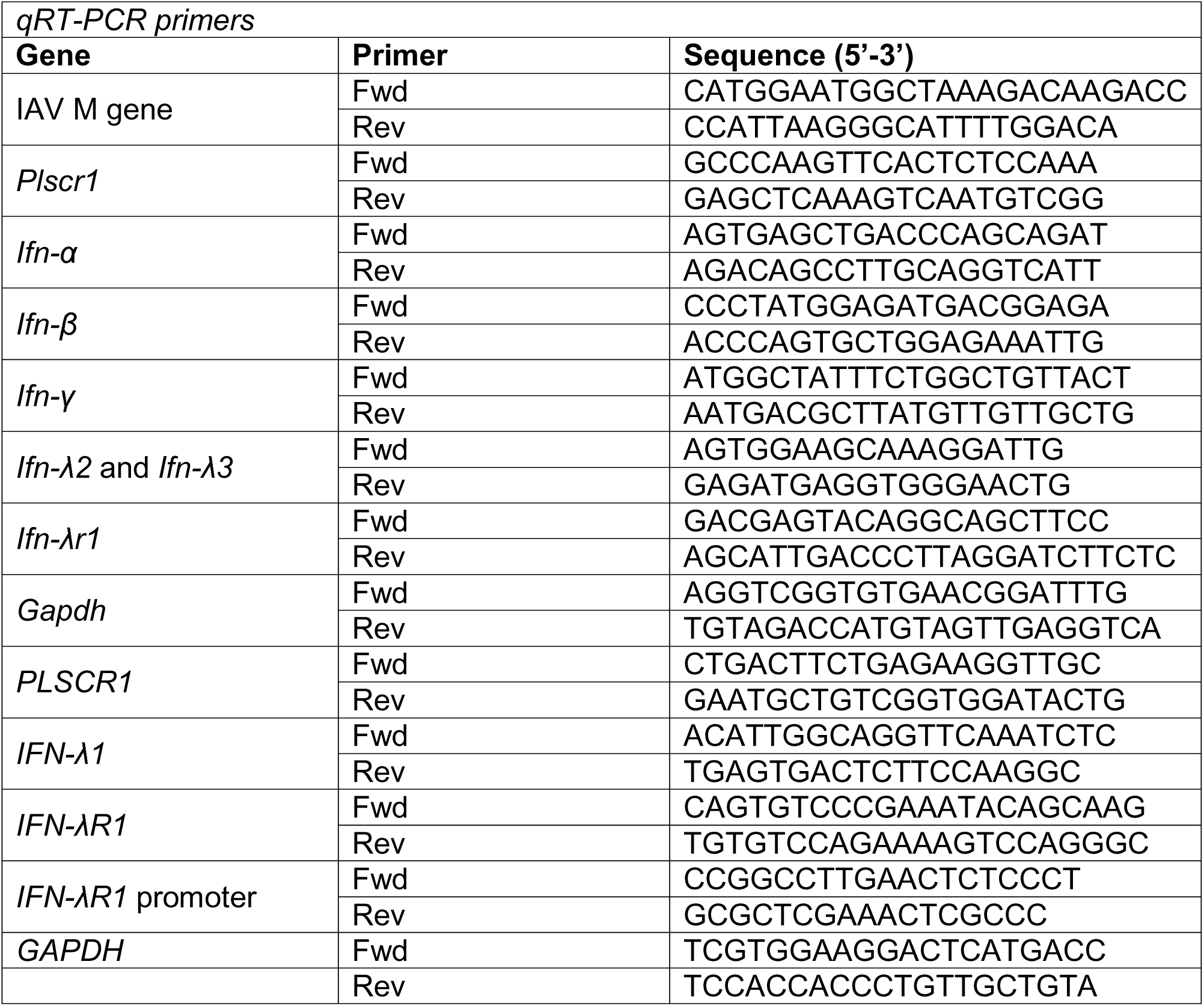
PCR Primer List.

### Immunohistology and immunofluorescence

For immunohistology, immediately after euthanasia, left lungs were inflated through the incision at the trachea with 10% buffered formalin and soaked in formalin at RT. They were then transferred to 70% ethanol prior to paraffin embedding, sectioning and staining with hematoxylin and eosin (H&E) performed at Brown University Molecular Pathology Core. Lungs were scanned with a VS200 slide scanner (Evident).

For immunofluorescence of mouse lungs, unstained paraffin-embedded lung sections were rehydrated in xylene (Fisher Scientific) and decreasing concentrations of ethanol (Pharmco). They were then steamed in antigen retrieval buffer (Abcam) for 30 minutes and blocked with 1% normal goat serum (Abcam) for 30 minutes at RT. Tissues were stained overnight at 4°C with goat anti-mouse Ifn-λr1 polyclonal antibody (1:300 dilution, Invitrogen, lot #WL3462592C), rabbit anti-mouse uteroglobin polyclonal antibody (1:300 dilution, Santa Cruz Biotechnology, lot #F0220), rabbit anti-mouse Spc polyclonal antibody (1:100 dilution, Santa Cruz Biotechnology, lot #C2007), mouse anti-mouse Foxj1 monoclonal antibody (1:300 dilution, Invitrogen, lot #2712325), and/or rabbit anti-mouse Plscr1 polyclonal antibody (1:300 dilution, Proteintech, no lot/clone #). Tissues were washed 3 times with PBS and stained with chicken anti-goat Alexa 594 (Invitrogen, lot #2318436), goat anti-rabbit Alexa 488 (Invitrogen, lot #2051237) and/or donkey anti-goat Alexa 405 (Invitrogen, lot #XI353690) for 1 hour at RT in the dark. For visualization of IAV, tissues were stained overnight at 4°C with goat anti-H1N1-FITC (1:500 dilution, US Biological, lot #L23091555 C23082906). After washing 3 times with PBS, sections were mounted with Vectashield® mounting medium with or without DAPI (Vector Laboratories). Lungs were visualized under an APEXVIEW APX100 microscope (Evident).

For immunofluorescence of cell cultures, cells were seeded in Millicell EZ slides (Millipore Sigma). After washing with ice-cold PBS, 4% paraformaldehyde (Boston BioProducts) was added on ice for 10 minutes for fixation and removed. If necessary, 0.2% Triton X-100 (Sigma-Aldrich) was added for 10 minutes to permeabilize the cells on ice, followed by 3 times of PBS wash. Cells were blocked with 5% bovine serum albumin (Fisher Scientifics) for 1 hour at RT. They were stained overnight at 4°C with mouse anti-human PLSCR1 monoclonal antibody (1:50 dilution, R&D Systems, clone #875327) and rabbit anti-human IFN-λR1 polyclonal antibody (1:100 dilution, Invitrogen, lot #ZB4228832A). Cells were washed 3 times with PBS and stained with chicken anti-mouse Alexa 594 (Invitrogen, lot #2482957) and goat anti-rabbit Alexa 488 for 1 hour at RT in the dark. After washing 3 times with PBS, slides were mounted with Vectashield® mounting medium with DAPI and visualized under a Nikon ECLIPSE Ti microscope.

When necessary, fluorescent images were analyzed using ImageJ[53]. Areas of interest were selected using freehand selection. Area, mean grey value and integrated density were measured. For H1N1 staining, uninfected lungs were used as background readings. For Ifn-λr1 staining, a region without fluorescence was used as a background reading. Corrected total cell fluorescence (CTCF) was calculated using the following formula: CTCF = Integrated Density – (Area of selection X Mean fluorescence of background readings).

### Proximity ligation assay (PLA)

Duolink® PLA fluorescence (Millipore Sigma) was performed according to manufacturer’s protocol. In brief, unstained paraffin-embedded lung sections were rehydrated, retrieved and blocked with Duolink® Blocking Solution for 60 minutes at 37°C. They were then incubated with goat anti-mouse Ifn-λr1 polyclonal antibody and rabbit anti-mouse Plscr1 polyclonal antibody overnight at 4°C. Next, slides were incubated with anti-goat PLUS and anti-rabbit MINUS probes for 1 hour at 37°C. For cell culture, A549 cells were seeded in Millicell EZ slides, fixed and permeabilized. After blocking, cells were incubated with rabbit anti-human IFN-λR1 polyclonal antibody and mouse anti-human PLSCR1 monoclonal antibody overnight at 4°C. Next, cells were incubated with anti-mouse PLUS and anti-rabbit MINUS probes for 1 hour at 37°C. Diluted ligase and polymerase were applied to samples for 30 minutes at 37°C in order. Finally, slides were mounted with Duolink® In Situ Mounting Medium with DAPI and imaged with an APEXVIEW APX100 microscope.

### IAV Infection and rhIFN-**λ**1 treatment in cell culture

Calu-3 cells at 80-90% confluency were washed twice with Opti-MEM (Gibco) and infected with IAV (WSN) at 0.1 multiplicity of infection (MOI) in Opti-MEM. One hour after viral attachment at 4°C, IAV supernatant was removed. Cells were washed once and incubated in fresh Opti-MEM for 23 hours at 37°C.

A549 cells at 80-90% confluency were washed once with PBS and infected with IAV (PR8) at 1, 5 or 10 MOI in PBS supplemented with BSA, CaCl_2_ and MgCl_2_. One hour after viral attainment at 37°C, IAV supernatant was removed. Cells were incubated in Opti-MEM with 10 μg/mL trypsin for 23 hours at 37°C.

Calu-3 or A549 cells were treated with 10 or 100 ng/mL of recombinant human IL-29/IFN-lambda 1 Protein (R&D Systems) for 6 or 24 hours at 37°C. In control Calu-3 groups, cells were first incubated with 1 μg/mL of anti-human interferon lambda receptor 1 neutralizing antibody (PBL Assay Science, clone #MMHLR-1) for 1 hour, and then treated with IFN-λ.

### Co-immunoprecipitation (Co-IP)

Cryopreserved mouse lungs were homogenized in Pierce RIPA buffer (Thermo Scientific). Protein concentrations were determined using Pierce BCA Protein Assay Kit (Thermo Scientific) according to the manufacturer’s protocol. Catch and Release® v2.0 Reversible Immunoprecipitation System (Millipore) was used according to manufacturer’s protocol. In brief, 500 μg of lung lysate, 2 μg of rabbit anti-human Plscr1 polyclonal antibody or negative control human IgG and 10 μL of antibody capture affinity ligand were added to the spin columns with resin slurry. The mixture was incubated on a rotator at 4°C overnight. The unbound flow-through was collected by centrifuge. Bound proteins were washed 3 times and eluted using denaturing elution buffer (5% SDS, 8M Urea, and 100mM Glycine).

### Western blot

Cryopreserved mouse lungs were homogenized in Pierce RIPA buffer. Cells in culture were washed with ice-cold PBS, lysed in Pierce RIPA buffer for 10 minutes on ice, and then scraped into RIPA buffer. Protein concentrations were determined using Pierce BCA Protein Assay Kit according to the manufacturer’s protocol. Twenty μg of each sample was loaded into 4-20% Mini-PROTEAN TGX Gels (Bio-Rad) and run for 1 hour 20 minutes at 125V in Tris/Glycine/SDS running buffer (Bio-Rad). Gels were transferred using Trans-Blot Turbo Transfer Pack (Bio-Rad). Membranes were incubated with rabbit anti-mouse/human Plscr1 polyclonal antibody (1:1000 dilution), rabbit anti-mouse/human Ifn-λr1 polyclonal antibody (1:1000 dilution, ABclonal, lot #0094910201), and/or mouse anti-mouse/human β-actin monoclonal antibody (1:2500 dilution, Santa Cruz Biotechnology, lot #F1323) overnight at 4°C. After washing with TBST (Boston BioProducts), membranes were incubated with HRP-linked anti-rabbit IgG (1:5000 dilution, Cell Signaling, lot #30) or HRP-linked anti-mouse IgG (1:5000 dilution, Cell Signaling, lot #36) for 1.5 hour at RT. Finally, membranes were treated with SuperSignal West Pico PLUS Chemiluminescent Substrate (Thermo Scientific) and imaged with ChemiDoc Imaging Systems (Bio-Rad).

### Chromatin-immunoprecipitation (CHIP)

SimpleCHIP® Enzymatic Chromatin IP (Cell Signaling) was performed according to manufacturer’s protocol. In brief, IAV-infected Calu-3 cells were fixed with formaldehyde to cross-link proteins to DNA. Next, chromatin was digested with Micrococcal Nuclease into 150-900 bp DNA/protein fragments. Nuclear membrane was broken by sonication. A portion of chromatin preparation was purified for DNA prior to immunoprecipitation to confirm the digestion by electrophoresis and concentration by OD260. One μg rabbit anti-mouse Plscr1 polyclonal antibody, negative control normal rabbit IgG, or positive control Histone H3 rabbit monoclonal antibody was added to 5μg chromatin preparation. The complex co-precipitated and was captured by Protein G magnetic beads. After 4 low salt washes and 2 high salt washes, chromatin was eluted and cross-links were reversed with proteinase K. DNA was purified using spin columns and analyzed by both standard PCR and qRT-PCR. For qRT-PCR, percent input method was used according to the following formula: Percent Input = 2% × 2^(C[T] 2%Input Sample − C[T] IP Sample)^.

### Transformation of competent cells

All *PLSCR1* plasmids were provided by Dr. John MacMicking’s Lab at Yale University. Plasmid DNA concentrations were determined using a spectrophotometer. Fifty ul of DH5α competent cells (Invitrogen) and 50ng of each plasmid was mixed and incubated for 30 minutes on ice. Then, the plasmid and competent cell mixtures were heat shocked for 45 seconds at 42°C, and cooled on ice for 2 minutes. Super Optimal broth with Catabolite repression (SOC) media (homemade with 2% tryptone, 0.5% yeast extract, 10mM NaCl, 2.5mM KCl, 10 mM MgCl2, 10 mM MgSO4, and 20 mM glucose) was added to bacteria. The cells were grown in a shacking incubator at 200RPM for 45 minutes at 37°C. The mixtures were then plated on LB agar ampicillin plates and incubated at 37°C overnight.

Single colonies were selected for each plate, inoculated in Super Optimal Broth (SOB) media (homemade with 2% tryptone, 0.5% yeast extract, 10mM NaCl, 2.5mM KCl, 10 mM MgCl2, and 10 mM MgSO4) and grown to OD600 of 0.6 at 37°C for 12-18 hours, with vigorous shacking of 200RPM. Cultures were then centrifuged at 6000G for 15 minutes at 4°C. Cell pallets were used for plasmid isolation.

### DNA plasmid isolation

DNA plasmids were isolated using the QIAfilter Maxi Cartridges and QIAGEN Plasmid Maxi Kit, according to the manufacture’s protocol. In brief, *PLSCR1*-transduced competent cell pellets were resuspended in buffer P1 containing RNase A solution and lyse blue. After adding Buffer P2, mixtures were incubated at RT for up to 5 minutes. Buffer P3 was then mixed with the lysates, transferred into the barrel of the QIAfilter cartridges and incubated at RT for 10 minutes. Cell lysates were filtered through the equilibrated QIAGEN-tip by gravity. DNA was eluted with Buffer QF and precipitated with isopropanol by centrifuging at 15000G for 10 minutes. Pellets were airdried for 10 minutes and dissolved in di-water. A spectrophotometer was used to determine plasmid concentrations.

### Lentiviral packaging

293T cells were grown in DMEM until 90-95% confluency and then switched to Opti-MEM for 2 hours. Ten ug of plasmid of interest was mixed with lipofectamine 300 transfection reagent (Invitrogen), pPACKH1 HIV Lentivector Packaging Kit (System Biosciences) and Opti-MEM, and incubated for 20mins at RT. After removing half media from the cell culture dishes, DNA-lipid complex was added and cells were incubated at 37°C with 5% CO2 for 6 hours. After 6 hours, culture supernatants were replaced with viral harvesting media (DMEM with glutamine, 10% FBS, 1% MEM non-essential amino acids, 1mM sodium pyruvate, 2% BSA, and 1:50 Hank’s solution) for overnight incubation. Viral supernatant was collected every day for the next 4 days by centrifuging cell culture supernatant at 1200G for 5 minutes. Viral supernatant was mixed with lentiviral concentrator solution (40% polyethylene glycol 8000 and 1.2M NaCl in PBS) at 4:1, and centrifuged at 1600G for 60 minutes at 4°C. The viral pellets were resuspended with Opti-MEM and stored at −80°C.

### Lentiviral transduction

*PLSCR1^-/-^* A549 cells were seeded in a 96 well plate with DMEM. Lentiviral stocks were serially diluted in DMEM and polybrene (Millipore Sigma) mixture from 1:10 to 1:10,000, and added to cell culture. The 96 well plate was centrifuged at 2000RPM for 30 minutes at RT and incubated at 37°C with 5% CO_2_ overnight. Media was replaced with fresh DMEM without polybrene the next day. After 48 hours post transduction, DMEM with 200ug/ml hygromycin was added for antibiotic selection for 10 days.

### Flow cytometry

Antibodies were titrated prior to confirming transduction efficiency by flow cytometry. One million cells were used for each sample. Samples were stained with LIVE/DEAD™ fixable violet dead cell stain for 405 nm excitation (1:1000, Invitrogen, lot #2581659) for 30 minutes. Surface and cytoplasm staining samples were fixed with 4% paraformaldehyde for 20 minutes at RT. Nuclear staining samples were fixed with true nuclear fix concentrate (BioLegend) for 45 minutes at RT. All samples were stained with mouse anti-human PLSCR1 monoclonal antibody (1:500) for 20 minutes, and chicken anti-mouse Alexa 594 (1:1000) for 20 minutes on ice. After fixation and each antibody incubation, surface staining samples were washed twice with FACS buffer (1% bovine serum albumin), cytoplasm staining samples were washed twice with Perm/Wash buffer (BD), and nuclear staining samples were washed twice with true nuclear perm buffer (BioLegend). Final samples were resuspended in FACS buffer. Flow cytometry analysis was performed with FACSAriaIIIu (BD) by Brown University Flow Cytometry and Sorting Facility.

### Cell coverage assay

After IAV infection, A549 cells in 12-well plates were fixed with 4% paraformaldehyde in PBS for 1 hour at RT. Cells were then stained with 1% crystal violet solution (Sigma-Aldrich) for 5 minutes, washed with di-water 3 times and air-dried. Plates were scanned with an APEXVIEW APX100 microscope. Brightfield images were converted to 8-bit and the threshold was adjusted using a dark background in ImageJ[53]. Areas of interest were selected using oval selection. Cell coverages were quantified by measuring mean grey values of each well.

### Plaque assay

For plaque assay of IAV titer in mouse lungs, a small piece of frozen lung was weighed and homogenized in 10mL decarbonated DMEM (MP Biomedicals) per 1g of lung. The homogenizer was sterilized in between samples with washes in the following order: twice with 0.1% SDS (Bio-Rad), 1:100 diluted bleach in di-water, once with 0.1% SDS, 0.001% Coomassie blue in di-water, twice with 70% ethanol, and once with sterile PBS. Homogenates were centrifuged at 2000G for 5 minutes at RT, and supernatant was collected.

For plaque assay of IAV titer in cell cultures, flu supernatant was collected, centrifuged at 2000G for 10 minutes, and filtered through 0.45μm filter.

MDCK cells were grown in 6-well plates to ∼80% confluency. Virus was diluted to desired concentrations with PBS supplemented with BSA, CaCl_2_ and MgCl_2_. Media was aspirated from MDCK cell cultures and replaced with 100μL of viral samples in each well. The plates were incubated for 1 hour at 37°C and agitated every 15 minutes to prevent from drying. After 1 hour, flu was aspirated and the wells were filled with 2mL plaque assay overlay (DMEM/F-12 (Gibco) supplemented with NaHCO_3_, BSA, DEAE dextran, and penicillin-streptomycin, mixed with 2% bacteriological agar (Oxoid) at 1:1 ratio). The plates were left at RT in the hood for 5 minutes to let the agar solidify, and then placed upside down in the 37°C incubator for 3 days until visible plaques were observed. Cells were fixed with 4% paraformaldehyde at RT for 1 hour. Agar overlay was removed with a metal spatula. Cells were stained with crystal violet solution gram (Harleco) for 5 minutes, and washed 3 times with di-water. Plates were air-dried. IAV titer was determined using the following formula: PFU/mL = # of plaques * 1/dilution factor * 10.

### Bulk RNA sequencing

Selected RNA isolated from mouse lungs in the same treatment group were pooled together into one sample. Pooled RNA concentrations were measured using a Nanodrop (Thermo Scientific). Quality control, library preparation, bulk RNA sequencing and data analysis was performed by Azenta Genomics/GENEWIZ. Heatmaps were made with expressions of selected genes in Morpheus (https://software.broadinstitute.org/morpheus). Hierarchical clustering was performed with one minus pearson correlation, complete linkage method and clustering by row. ISGs were identified using the Interferome database, with selections of a fold change ≥2 in all cell types in lung of *Mus musculus*[25].

### Single cell RNA sequencing analysis

Single cell RNA sequencing dataset were generated from uninfected or IAV-infected mouse lungs at 0, 1, 3, 6 and 21 dpi as previously described[28]. Single-cell transcriptomic libraries were generated using the 5′ Gene Expression Kit (V2, 10X Genomics) according to the manufacturer’s instructions with the addition of primers to amplify HTOs during cDNA amplification[28]. Sequencing was performed on the Illumina NovaSeq to generate approximately 500 million reads per sample[28]. The 10X gene-expression data were processed and normalized using CellRanger (v.3.0.2, 10X Genomics)[28]. QC was performed first by excluding any gene that was not present in at least 0.1% of total called cells and then by excluding cells that exhibited extremes in the species-specific distributions of: the number of genes expressed (<100 or >5,550), the number of mRNA molecules (>30,000), or the percentage of expression owed to mitochondrial genes (>7.5%)[28]. Scaling, principal component analysis, dimensionality reduction and cell clustering were performed using Seurat algorithm as previously described[28, 54]. Cell clusters were annotated using known markers from the literature (Table S2).

All following analysis were performed using R Statistical Software (v4.3.0)[55]. Two-dimensional UMAPs were generated with DimPlot (Seurat)[54]. Bar charts of proportion and cell count of each cluster were plotted with geom_col (ggplot2)[56]. Violin plots of *Plscr1* expression in different cell clusters were plotted with VlnPlot (Seurat)[54]. Differentially expressed genes for ciliated epithelial cell clusters at different timepoints in comparison to 0 dpi were identified with FindMarkers (Seurat)[54]. Those genes were used for Gene Ontology (GO) analysis with enrichGo (clusterProfiler)[57], and heatmaps with DoHeatmap (Seurat)[54].

**Table S2.** scRNA-Seq Cluster Annotations.

| *scRNA-seq cluster annotations* |  |  |
| --- | --- | --- |
| **Cluster** | **Cell Type** | **Transcriptional Marker** |
| 0 | AT2 Cells | *Epcam, Sftpc* |
| 1 | Mfap5+ Fibroblasts | *Pdgfra, Mfap5* |
| 2 | Microvascular Endothelial Cells | *Pecam1, Gpihbp1* |
| 3 | Damage-Responsive Fibroblasts | *Pdgfra, Wt1* |
| 4 | T Cells | *Ptprc, Cd3e* |
| 5 | Activated Microvascular Endothelial Cells | *Pecam1, Gpihbp1, S1pr1* |
| 6 | Airway Smooth Muscle Cells | *Hhip, Acta2* |
| 7 | IAV-Infected Epithelial Cells | *Epcam, Flu* |
| 8 | Microvascular Endothelial Cells | *Pecam1, Gpihbp1* |
| 9 | High Mitochondrial Content Cells |  |
| 10 | Krt8+ Epithelial Cells | *Epcam, Krt8* |
| 11 | AT2 Cells | *Epcam, Sftpc* |
| 12 | Ciliated Epithelial Cells | *Epcam, Foxj* |
| 13 | Club Cells | *Epcam, Scgb3a2* |
| 14 | B Cells | *Ptprc, Cd19* |
| 15 | Myofibroblasts | *Pdgfra, Acta2* |
| 16 | Neutrophils | *Ptprc, Mmp9* |
| 17 | Alveolar Fibroblasts | *Pdgfra, Wnt2* |
| 18 | Epcam+Pecam1+ Cells | *Epcam, Pecam1* |
| 19 | Monocytes | *Ptprc, Ly6c2* |
| 20 | Interferon-Responsive Fibroblasts | *Pdgfra, Bst2* |
| 21 | Mesothelial Cells | *Msln, Wt1* |
| 22 | Epcam+Col1a2+ EMT Cells | *Epcam, Col1a2* |
| 23 | Interstitial Macrophages | *Ptprc, Cd68, C1qb* |
| 24 | Lymphatic Endothelial Cells | *Pecam1, Prox1* |
| 25 | Aerocytes | *Pecam 1, Car4* |
| 26 | Vascular Smooth Muscle Cells | *Acta2, Pdgfrb, Notch3* |
| 27 | AT1 Cells | *Epcam, Akap5, Ager* |
| 28 | Macrovascular Endothelial Cells | *Pecam1, Vwf* |
| 29 | Ciliated Epithelial Cells | *Epcam, Foxj1* |
| 30 | Alveolar Macrophages | *Ptprc, Cd68, Chil3* |
| 31 | NK Cells | *Ptprc, Klrb1c* |
| 32 | Epcam+Col1a2+ EMT Cells | *Epcam, Col1a2* |
| 33 | Immune Doublets | *Ptprc, Cd3e, Cd19* |
| 34 | Regulatory T cells | *Cd4, Il2ra* |
| 35 | Basophils/Mast Cells | *Ptprc, Fcer1a, Gata2* |
| 36 | Multiplets |  |
| 37 | Platelets |  |

### Quantitation and statistical analysis

Data were analyzed on GraphPad Prism software. Logrank (Mantel-Cox) test was used to compare survival rates. Ordinary two-way ANOVA tests were used to compare weight losses. Statistical significance of differences was assessed using the parametric Student’s two-tailed t tests for all other normally distributed data. Differences were considered significant when p < 0.05. Outlier tests with the ROUT method (Q = 1%) were used to identify and remove any outliers.

## ACKNOWLEDGEMENTS

The authors thank A. Ayala, A. Jamieson and C. Lee for advice; J. MacMicking for sharing *PLSCR1^-/-^* and *T16F^-/-^* A549 cells and all *PLSCR1* plasmids; Brown University Molecular Pathology Core for histopathology; and Brown University Flow Cytometry and Sorting Facility for flow cytometry.

This work was supported by grants R01 HL146498 (YZ), P20 GM103652 (YZ), U54 GM115677 (YZ), T32 HL134625 (PS), AHA 24TPA1277918 (YZ), ATS 23-24PHP11 (YZ), R01 AI183155 (SL), and R35 HL145242 (MJH).

During the preparation of this work the author(s) used ChatGPT developed by OpenAI in order to check grammar and improve readability. After using this tool/service, the author(s) reviewed and edited the content as needed and take(s) full responsibility for the content of the publication.

---

**Supplemental Figure 1. Heatmap of Differential Expressions of All ISGs in Whole Lungs by RNA-seq.**

Gene expressions were compared between groups for each row and color-labeled from row minimum (blue) to row maximum (red). Enlarged heatmaps of Figure 3C.

**Supplemental Figure 2. Requirement of *Plscr1* in IFN-λ Signaling Independent of Viral Titer**

*Wt* and *Plscr1^-/-^* mice were intranasally given 2.5 μg/g of body weight of poly(I:C) (HMW) constitutively for 6 days and sacrificed on day 7.

(A) Scheme of experiment.

(B) Total BAL leukocyte numbers.

(C) Differential cell counts in BAL.

(D, F-G) Whole lungs were analyzed for *Ifn-*α, *Ifn-*β, *Ifn-*γ, *Ifn-*λ (D); *Plscr1* (F); and *Ifn-* λ*r1* (G) RNA by qRT-PCR.

(A) (E) Representative lung sections stained with H&E. Scale bars represent 3 mm (main) and 200 μm (inlays).

Data are expressed as mean ± SEM of n = 5-12 mice/group. All data were pooled from three independent experiments. ns, not significant, *p < 0.05, ***p<0.001.

**Supplemental Figure 3. Co-Immunoprecipitation of Plscr1 and Ifn-λr1 in Whole Mouse Lungs Followed by Western Blot.**

Whole gel images of Figure 4A.

**Supplemental Figure 4. PLSCR1 Transduction Efficiency and Distribution**

*PLSCR1* plasmids on PLV-EF1a-IRES-Hygro backbone were packaged into GFP-expressing lentivirus. *PLSCR1^-/-^*A549 cells were transduced using lentivirus. After a 10-day hygromycin selection, cells were analyzed using flow cytometry.

(A) Gating strategy for live, GFP+ and PLSCR1+ A549 cells.

(B) Surface, cytoplasm and nuclear expression of PLSCR1. Data are presented as fold change compared to *WT-PLSCR1^-/-^* A549 cells.

(C) Lentiviral transduction efficiency.

**Supplemental Figure 5. The Relative Contribution of the Type 3 IFN Pathway to Plscr1-Mediated Antiviral Immunity.**

*Wt*, *Plscr1^-/-^*, *Ifn-*λ*r1^-/-^* and *Plscr1^-/-^Ifn-*λ*r1^-/-^* mice were exposed to sublethal (300 pfu) IAV (WSN) infection and sacrificed at 3 dpi.

(A) Representative immunofluorescent staining for DAPI and Ifn-λr1 in lungs.

(B) Mean relative weight of mice.

(C) Total BAL leukocyte numbers.

(D) Neutrophil percentages in BAL.

(E) Infectious viral titer in the lungs was assessed by plaque assays.

**Supplemental Figure 6. Proportion and Cell Count of Each Cluster in Single-Cell RNA Sequencing.**

Wt mice were exposed to 2500 EID50 IAV (PR8) infection. Lungs were used for single-cell RNA sequencing analysis at 0, 1, 3, 6 and 21 dpi.

(A) Proportion of each cluster.

(B) Cell count of each cluster.

**Supplemental Figure 7. Time-Dependent *Plscr1* Expressions in All Epithelial Cell Clusters Other than Ciliated Epithelial Cells.**

*(A-K) Wt* mice were exposed to 2500 EID50 IAV (PR8) infection. Lungs were used for single-cell RNA sequencing analysis at 0, 1, 3, 6 and 21 dpi.

**Supplemental Figure 8. Unaffected Susceptibility of *Plscr1^floxStop^LysM-Cre^+^* Mice to Influenza Virus Infection**

*Plscr1^floxStop^*and *Plscr1^floxStop^LysM-Cre^+^*mice were exposed to sublethal (300 pfu) IAV (WSN) infection.

(A) Validation of *Plscr1* overexpression in lungs of *Plscr1^floxStop^LysM-Cre^+^* mice by qRT-PCR.

(B) Mean relative weight of mice.

(C) Total BAL leukocyte numbers.

(D) Differential cell counts in BAL.

(E) Viral RNA load in the lungs was assessed by quantifying M gene by qRT-PCR.

(F) Representative lung sections stained with H&E.

(G-I) Whole lungs were analyzed for *Ifn-*α, *Ifn-*β, *Ifn-*γ, *Ifn-*λ (G); *Ifn-*λ*r1* (H); and *Plscr1* (I) RNA by qRT-PCR.

Data are expressed as mean ± SEM of n = 15-16 mice/group for weight loss. For the rest analysis, n= 3-7 mice/group. All data were pooled from three independent experiments. ns, not significant, *p < 0.05, **p < 0.01. dpi, days post infection.

