## Supplemental Figures for "Phospholipid Scramblase 1 (PLSCR1) Regulates Interferon-Lambda Receptor 1 (IFN-λR1) and IFN-λ Signaling in Influenza A Virus (IAV) Infection"

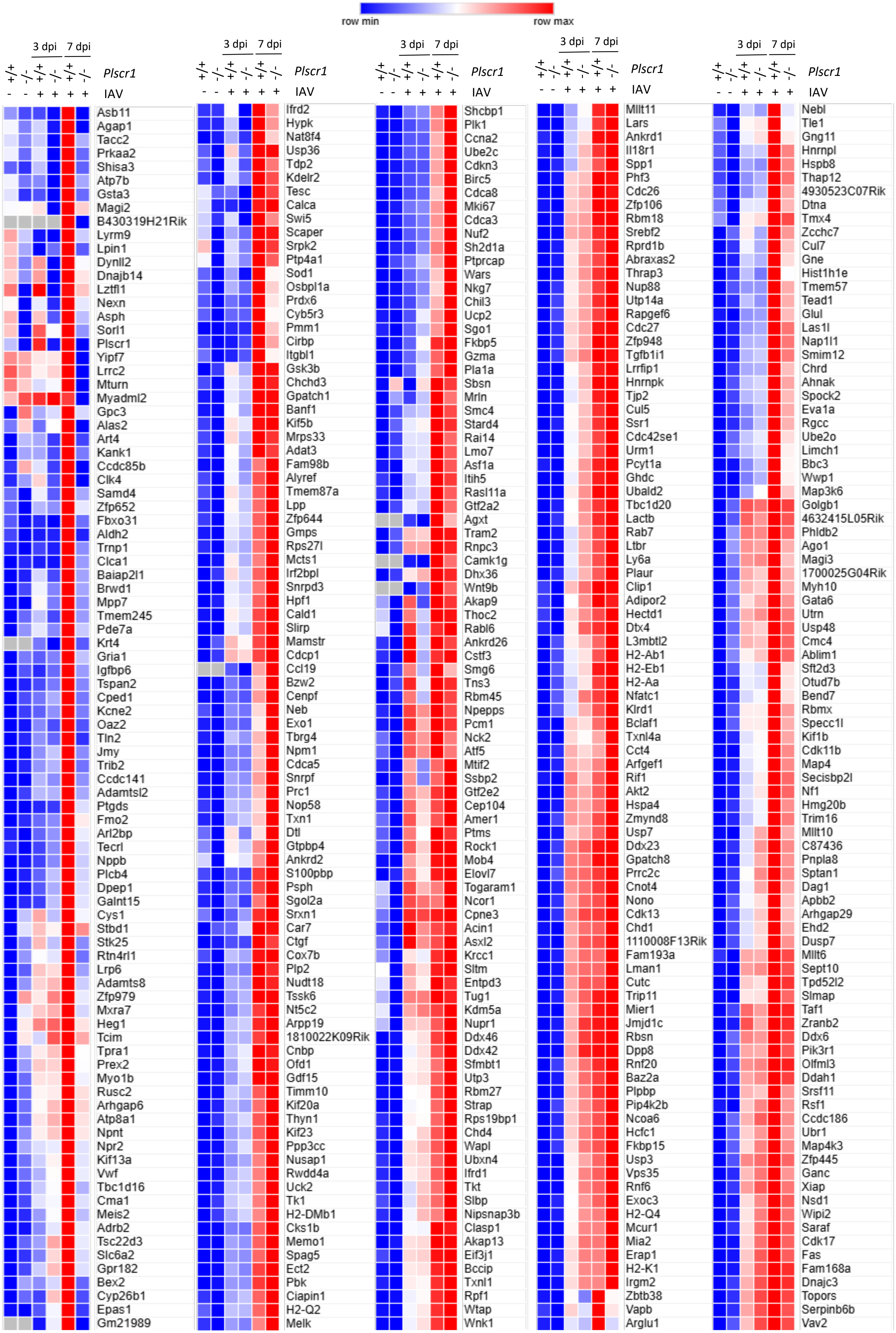

row min
row max
3 dpi
7 dpi
+/+ -/- +/+ -/- +/+ -/- Plscr1
- - + + + + IAV
Asb11
Agap1
Tacc2
Prkaa2
Shisa3
Atp7b
Gsta3
Magi2
B430319H21Rik
Lyrm9
Lpin1
Dynll2
Dnajb14
Lztfl1
Nexn
Asph
Sorl1
Plscr1
Yipf7
Lrrc2
Mturn
Myadml2
Gpc3
Alas2
Art4
Kank1
Ccdc85b
Clk4
Samd4
Zfp652
Fbxo31
Aldh2
Trnp1
Clca1
Baiap2l1
Brwd1
Mpp7
Tmem245
Pde7a
Krt4
Gria1
Igfbp6
Tspan2
Cped1
Kcne2
Oaz2
Tln2
Jmy
Trib2
Ccdc141
Adamtsl2
Ptgds
Fmo2
Arl2bp
Tecrl
Nppb
Plcb4
Dpep1
Galnt15
Cys1
Stbd1
Stk25
Rtn4rl1
Lrp6
Adamts8
Zfp979
Mxra7
Heg1
Tcim
Tpra1
Prex2
Myo1b
Rusc2
Arhgap6
Atp8a1
Npnt
Npr2
Kif13a
Vwf
Tbc1d16
Cma1
Meis2
Adrb2
Tsc22d3
Slc6a2
Gpr182
Bex2
Cyp26b1
Epas1
Gm21989
3 dpi
7 dpi
+/+ -/- +/+ -/- +/+ -/- Plscr1
- - + + + + IAV
Ifrd2
Hypk
Nat8f4
Usp36
Tdp2
Kdelr2
Tesc
Calca
Swi5
Scaper
Srpk2
Ptp4a1
Sod1
Osbpl1a
Prdx6
Cyb5r3
Pmm1
Cirbp
Itgbl1
Gsk3b
Chchd3
Gpatch1
Banf1
Kif5b
Mrps33
Adat3
Fam98b
Alyref
Tmem87a
Lpp
Zfp644
Gmps
Rps27l
Mcts1
Irf2bpl
Snrpd3
Hpf1
Cald1
Slirp
Mamstr
Cdcp1
Ccl19
Bzw2
Cenpf
Neb
Exo1
Tbrg4
Npm1
Cdca5
Snrpf
Prc1
Nop58
Txn1
Dtl
Gtpbp4
Ankrd2
S100pbp
Psph
Sgol2a
Srxn1
Car7
Ctgf
Cox7b
Plp2
Nudt18
Tssk6
Nt5c2
Arpp19
1810022K09Rik
Cnbp
Ofd1
Gdf15
Timm10
Kif20a
Thyn1
Kif23
Ppp3cc
Nusap1
Rwdd4a
Uck2
Tk1
H2-DMb1
Cks1b
Memo1
Spag5
Ect2
Pbk
Ciapin1
H2-Q2
Melk
3 dpi
7 dpi
+/+ -/- +/+ -/- +/+ -/- Plscr1
- - + + + + IAV
Shcbp1
Plk1
Ccna2
Ube2c
Cdkn3
Birc5
Cdca8
Mki67
Cdca3
Nuf2
Sh2d1a
Ptprcap
Wars
Nkg7
Chil3
Ucp2
Sgo1
Fkbp5
Gzma
Pla1a
Sbsn
Mrln
Smc4
Stard4
Rai14
Lmo7
Asf1a
Itih5
Rasl11a
Gtf2a2
Agxt
Tram2
Rnpc3
Camk1g
Dhx36
Wnt9b
Akap9
Thoc2
Rabl6
Ankrd26
Cstf3
Smg6
Tns3
Rbm45
Npepps
Pcm1
Nck2
Atf5
Mtif2
Ssbp2
Gtf2e2
Cep104
Amer1
Ptms
Rock1
Mob4
Elovl7
Togaram1
Ncor1
Cpne3
Acin1
Asxl2
Krcc1
Sltm
Entpd3
Tug1
Kdm5a
Nupr1
Ddx46
Ddx42
Sfmbt1
Utp3
Rbm27
Strap
Rps19bp1
Chd4
Wapl
Ubxn4
Ifrd1
Tkt
Slbp
Nipsnap3b
Clasp1
Akap13
Eif3j1
Bccip
Txnl1
Rpf1
Wtap
Wnk1
3 dpi
7 dpi
+/+ -/- +/+ -/- +/+ -/- Plscr1
- - + + + + IAV
Mllt11
Lars
Ankrd1
Il18r1
Spp1
Phf3
Cdc26
Zfp106
Rbm18
Srebf2
Rprd1b
Abraxas2
Thrap3
Nup88
Utp14a
Rapgef6
Cdc27
Zfp948
Tgfb1i1
Lrrfip1
Hnrnpk
Tjp2
Cul5
Ssr1
Cdc42se1
Urm1
Pcyt1a
Ghdc
Ubald2
Tbc1d20
Lactb
Rab7
Ltbr
Ly6a
Plaur
Clip1
Adipor2
Hectd1
Dtx4
L3mbtl2
H2-Ab1
H2-Eb1
H2-Aa
Nfatc1
Klrd1
Bclaf1
Txnl4a
Cct4
Arfgef1
Rif1
Akt2
Hspa4
Zmynd8
Usp7
Ddx23
Gpatch8
Prrc2c
Cnot4
Nono
Cdk13
Chd1
1110008F13Rik
Fam193a
Lman1
Cutc
Trip11
Mier1
Jmjd1c
Rbsn
Dpp8
Rnf20
Baz2a
Plpbp
Pip4k2b
Ncoa6
Hcfc1
Fkbp15
Usp3
Vps35
Rnf6
Exoc3
H2-Q4
Mcur1
Mia2
Erap1
H2-K1
Irgm2
Zbtb38
Vapb
Arglu1
3 dpi
7 dpi
+/+ -/- +/+ -/- +/+ -/- Plscr1
- - + + + + IAV
Nebl
Tle1
Gng11
Hnrnpl
Hspb8
Thap12
4930523C07Rik
Dtna
Tmx4
Zcchc7
Cul7
Gne
Hist1h1e
Tmem57
Tead1
Glul
Las1l
Nap1l1
Smim12
Chrd
Ahnak
Spock2
Eva1a
Rgcc
Ube2o
Limch1
Bbc3
Wwp1
Map3k6
Golgb1
4632415L05Rik
Phldb2
Ago1
Magi3
1700025G04Rik
Myh10
Gata6
Utrn
Usp48
Cmc4
Ablim1
Sft2d3
Otud7b
Bend7
Rbmx
Specc1l
Kif1b
Cdk11b
Map4
Secisbp2l
Nf1
Hmg20b
Trim16
Mllt10
C87436
Pnpla8
Sptan1
Dag1
Apbb2
Arhgap29
Ehd2
Dusp7
Mllt6
Sept10
Tpd52l2
Slmap
Taf1
Zranb2
Ddx6
Pik3r1
Olfml3
Ddah1
Srsf11
Rsf1
Ccdc186
Ubr1
Map4k3
Zfp445
Ganc
Xiap
Nsd1
Wipi2
Saraf
Cdk17
Fas
Fam168a
Dnajc3
Topors
Serpinb6b
Vav2

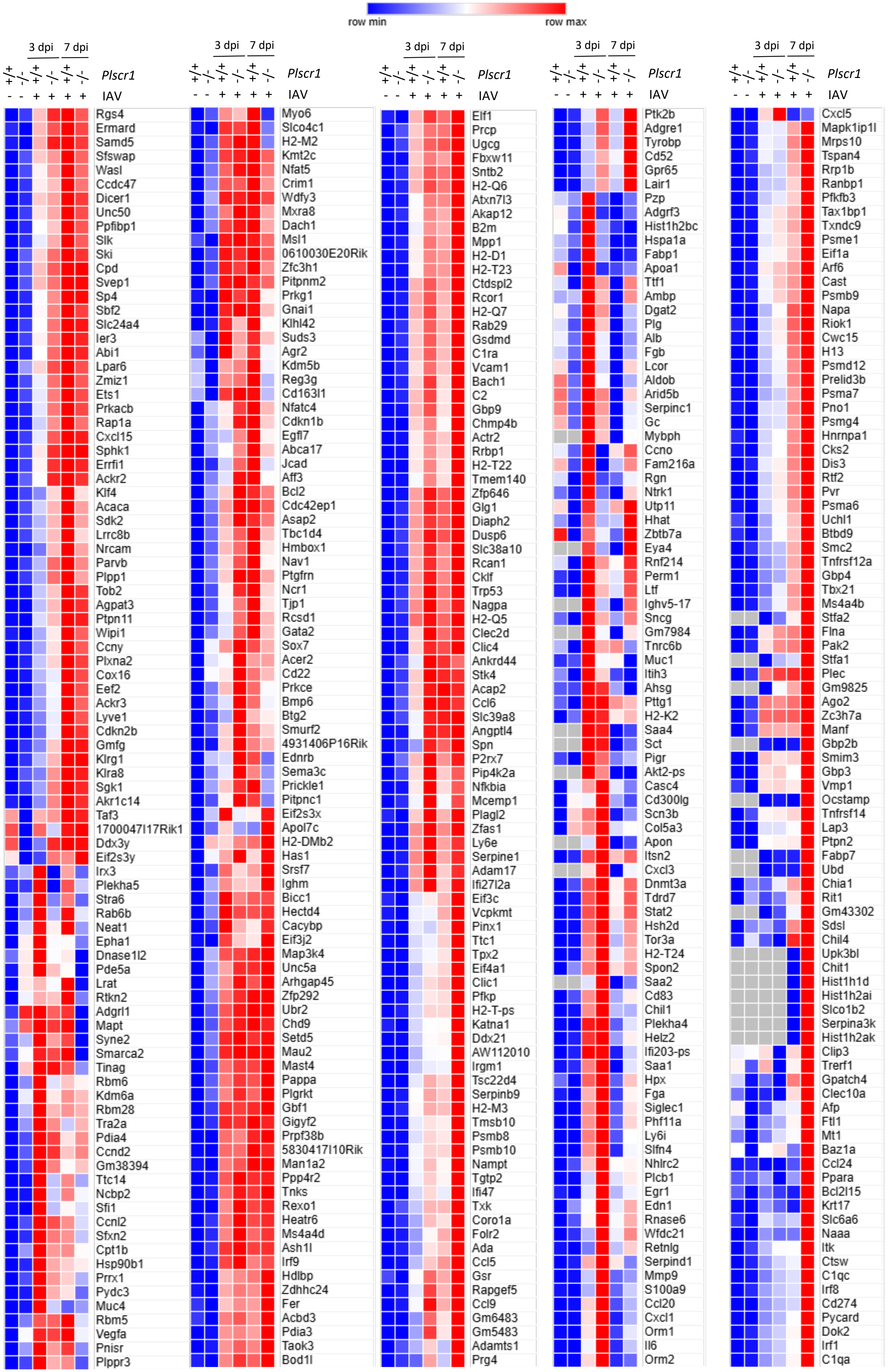
row min
row max
3 dpi
7 dpi
+/+ -/- +/+ -/- +/+ -/-
Plscr1
- - + + + +
IAV
Rgs4
Ermard
Samd5
Sfswap
Wasl
Ccdc47
Dicer1
Unc50
Ppfibp1
Slk
Ski
Cpd
Svep1
Sp4
Sbf2
Slc24a4
Ier3
Abi1
Lpar6
Zmiz1
Ets1
Prkacb
Rap1a
Cxcl15
Sphk1
Errfi1
Ackr2
Klf4
Acaca
Sdk2
Lrrc8b
Nrcam
Parvb
Plpp1
Tob2
Agpat3
Ptpn11
Wipi1
Ccny
Plxna2
Cox16
Eef2
Ackr3
Lyve1
Cdkn2b
Gmfg
Klrg1
Klra8
Sgk1
Akr1c14
Taf3
1700047I17Rik1
Ddx3y
Eif2s3y
Irx3
Plekha5
Stra6
Rab6b
Neat1
Epha1
Dnase1l2
Pde5a
Lrat
Rtkn2
Adgrl1
Mapt
Syne2
Smarca2
Tinag
Rbm6
Kdm6a
Rbm28
Tra2a
Pdia4
Ccnd2
Gm38394
Ttc14
Ncbp2
Sfi1
Ccnl2
Sfxn2
Cpt1b
Hsp90b1
Prrx1
Pydc3
Muc4
Rbm5
Vegfa
Pnisr
Plppr3
Myo6
Slco4c1
H2-M2
Kmt2c
Nfat5
Crim1
Wdfy3
Mxra8
Dach1
Msl1
0610030E20Rik
Zfc3h1
Pitpnm2
Prkg1
Gnai1
Klhl42
Suds3
Agr2
Kdm5b
Reg3g
Cd163l1
Nfatc4
Cdkn1b
Egfl7
Abca17
Jcad
Aff3
Bcl2
Cdc42ep1
Asap2
Tbc1d4
Hmbox1
Nav1
Ptgfrn
Ncr1
Tjp1
Rcsd1
Gata2
Sox7
Acer2
Cd22
Prkce
Bmp6
Btg2
Smurf2
4931406P16Rik
Ednrb
Sema3c
Prickle1
Pitpnc1
Eif2s3x
Apol7c
H2-DMb2
Has1
Srsf7
Ighm
Bicc1
Hectd4
Cacybp
Eif3j2
Map3k4
Unc5a
Arhgap45
Zfp292
Ubr2
Chd9
Setd5
Mau2
Mast4
Pappa
Plgrkt
Gbf1
Gigyf2
Prpf38b
5830417I10Rik
Man1a2
Ppp4r2
Tnks
Rexo1
Heatr6
Ms4a4d
Ash1l
Irf9
Hdlbp
Zdhhc24
Fer
Acbd3
Pdia3
Taok3
Bod1l
Elf1
Prcp
Ugcg
Fbxw11
Sntb2
H2-Q6
Atxn7l3
Akap12
B2m
Mpp1
H2-D1
H2-T23
Ctdspl2
Rcor1
H2-Q7
Rab29
Gsdmd
C1ra
Vcam1
Bach1
C2
Gbp9
Chmp4b
Actr2
Rrbp1
H2-T22
Tmem140
Zfp646
Glg1
Diaph2
Dusp6
Slc38a10
Rcan1
Cklf
Trp53
Nagpa
H2-Q5
Clec2d
Clic4
Ankrd44
Stk4
Acap2
Ccl6
Slc39a8
Angptl4
Spn
P2rx7
Pip4k2a
Nfkbia
Mcemp1
Plagl2
Zfas1
Ly6e
Serpine1
Adam17
Ifi27l2a
Eif3c
Vcpkmt
Pinx1
Ttc1
Tpx2
Eif4a1
Clic1
Pfkp
H2-T-ps
Katna1
Ddx21
AW112010
Irgm1
Tsc22d4
Serpinb9
H2-M3
Tmsb10
Psmb8
Psmb10
Nampt
Tgtp2
Ifi47
Txk
Coro1a
Folr2
Ada
Ccl5
Gsr
Rapgef5
Ccl9
Gm6483
Gm5483
Adamts1
Prg4
Ptk2b
Adgre1
Tyrobp
Cd52
Gpr65
Lair1
Pzp
Adgrf3
Hist1h2bc
Hspa1a
Fabp1
Apoa1
Ttf1
Ambp
Dgat2
Plg
Alb
Fgb
Lcor
Aldob
Arid5b
Serpinc1
Gc
Mybph
Ccno
Fam216a
Rgn
Ntrk1
Utp11
Hhat
Zbtb7a
Eya4
Rnf214
Perm1
Ltf
Ighv5-17
Sncg
Gm7984
Tnrc6b
Muc1
Itih3
Ahsg
Pttg1
H2-K2
Saa4
Sct
Pigr
Akt2-ps
Casc4
Cd300lg
Scn3b
Col5a3
Apon
Itsn2
Cxcl3
Dnmt3a
Tdrd7
Stat2
Hsh2d
Tor3a
H2-T24
Spon2
Saa2
Cd83
Chil1
Plekha4
Helz2
Ifi203-ps
Saa1
Hpx
Fga
Siglec1
Phf11a
Ly6i
Slfn4
Nhlrc2
Plcb1
Egr1
Edn1
Rnase6
Wfdc21
Retnlg
Serpind1
Mmp9
S100a9
Ccl20
Cxcl1
Orm1
Il6
Orm2
Cxcl5
Mapk1ip1l
Mrps10
Tspan4
Rrp1b
Ranbp1
Pfkfb3
Tax1bp1
Txndc9
Psme1
Eif1a
Arf6
Cast
Psmb9
Napa
Riok1
Cwc15
H13
Psmd12
Prelid3b
Psma7
Pno1
Psmg4
Hnrnpa1
Cks2
Dis3
Rtf2
Pvr
Psma6
Uchl1
Btbd9
Smc2
Tnfrsf12a
Gbp4
Tbx21
Ms4a4b
Stfa2
Flna
Pak2
Stfa1
Plec
Gm9825
Ago2
Zc3h7a
Manf
Gbp2b
Smim3
Gbp3
Vmp1
Ocstamp
Tnfrsf14
Lap3
Ptpn2
Fabp7
Ubd
Chia1
Rit1
Gm43302
Sdsl
Chil4
Upk3bl
Chit1
Hist1h1d
Hist1h2ai
Slco1b2
Serpina3k
Hist1h2ak
Clip3
Trerf1
Gpatch4
Clec10a
Afp
Ftl1
Mt1
Baz1a
Ccl24
Ppara
Bcl2l15
Krt17
Slc6a6
Naaa
Itk
Ctsw
C1qc
Irf8
Cd274
Pycard
Dok2
Irf1
C1qa

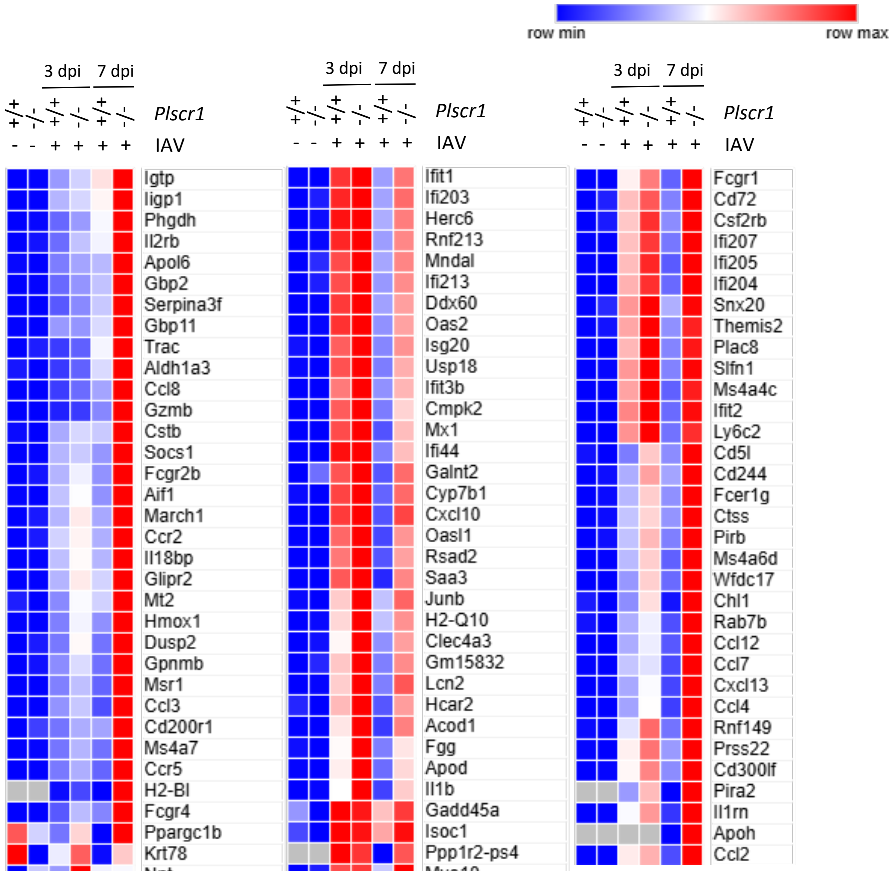

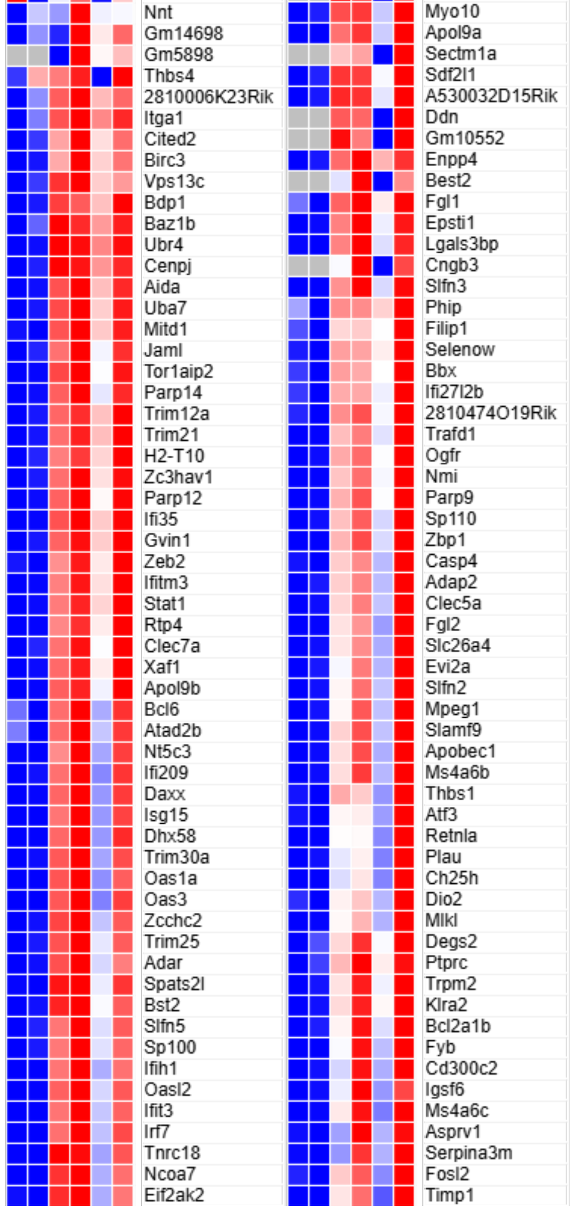


**Supplemental Figure 1. Heatmap of Differential Expressions of All ISGs in Whole Lungs by RNA-seq.** Gene expressions were compared between groups for each row and color-labeled from row minimum (blue) to row maximum (red). Enlarged heatmaps of Figure 3C.

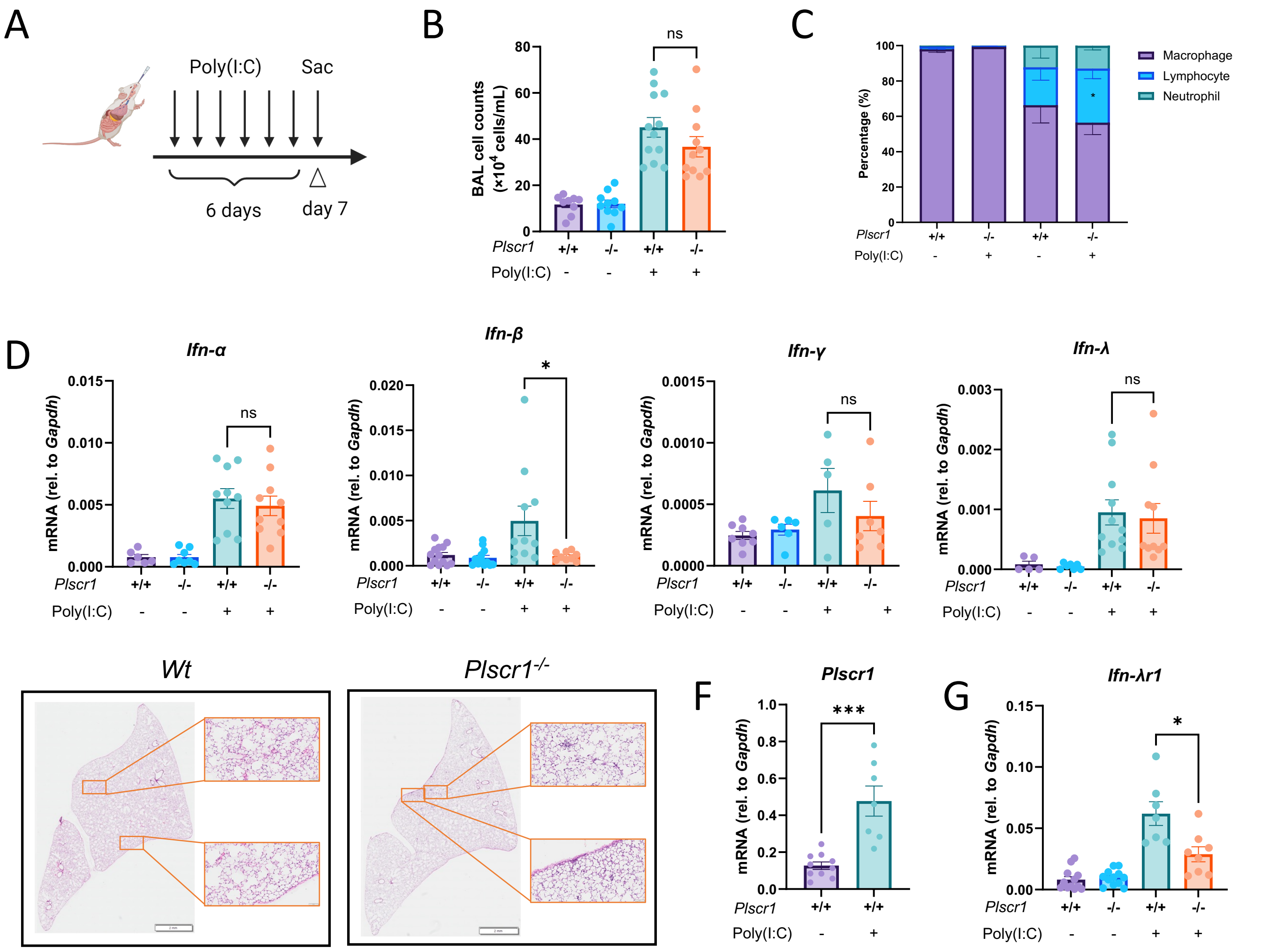


**Supplemental Figure 2. Requirement of *Plscr1* in IFN-λ Signaling Independent of Viral Titer**

*Wt* and *Plscr1*$^{-/-}$ mice were intranasally given 2.5 µg/g of body weight of poly(I:C) (HMW) constitutively for 6 days and sacrificed on day 7.

(A) Scheme of experiment.

(B) Total BAL leukocyte numbers.

(C) Differential cell counts in BAL.

(D, F-G) Whole lungs were analyzed for *Ifn-α*, *Ifn-β*, *Ifn-γ*, *Ifn-λ* (D); *Plscr1* (F); and *Ifn-λr1* (G) RNA by qRT-PCR.

(E) Representative lung sections stained with H&E. Scale bars represent 3 mm (main) and 200 µm (inlays).

Data are expressed as mean ± SEM of n = 5-12 mice/group. All data were pooled from three independent experiments. ns, not significant, *p < 0.05, ***p<0.001.

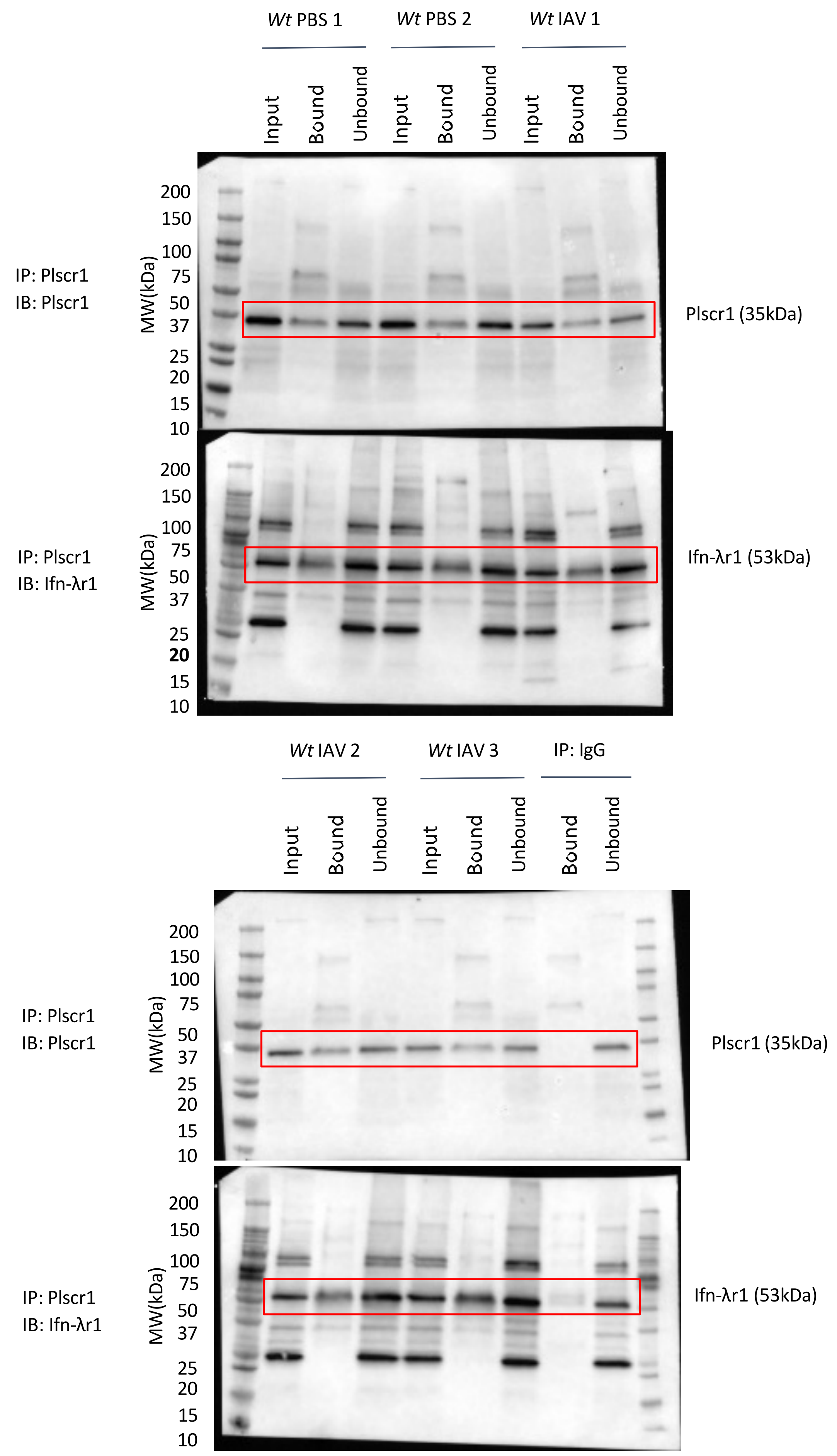


**Supplemental Figure 3. Co-Immunoprecipitation of Plscr1 and Ifn-λr1 in Whole Mouse Lungs Followed by Western Blot.**
Whole gel images of Figure 4A.

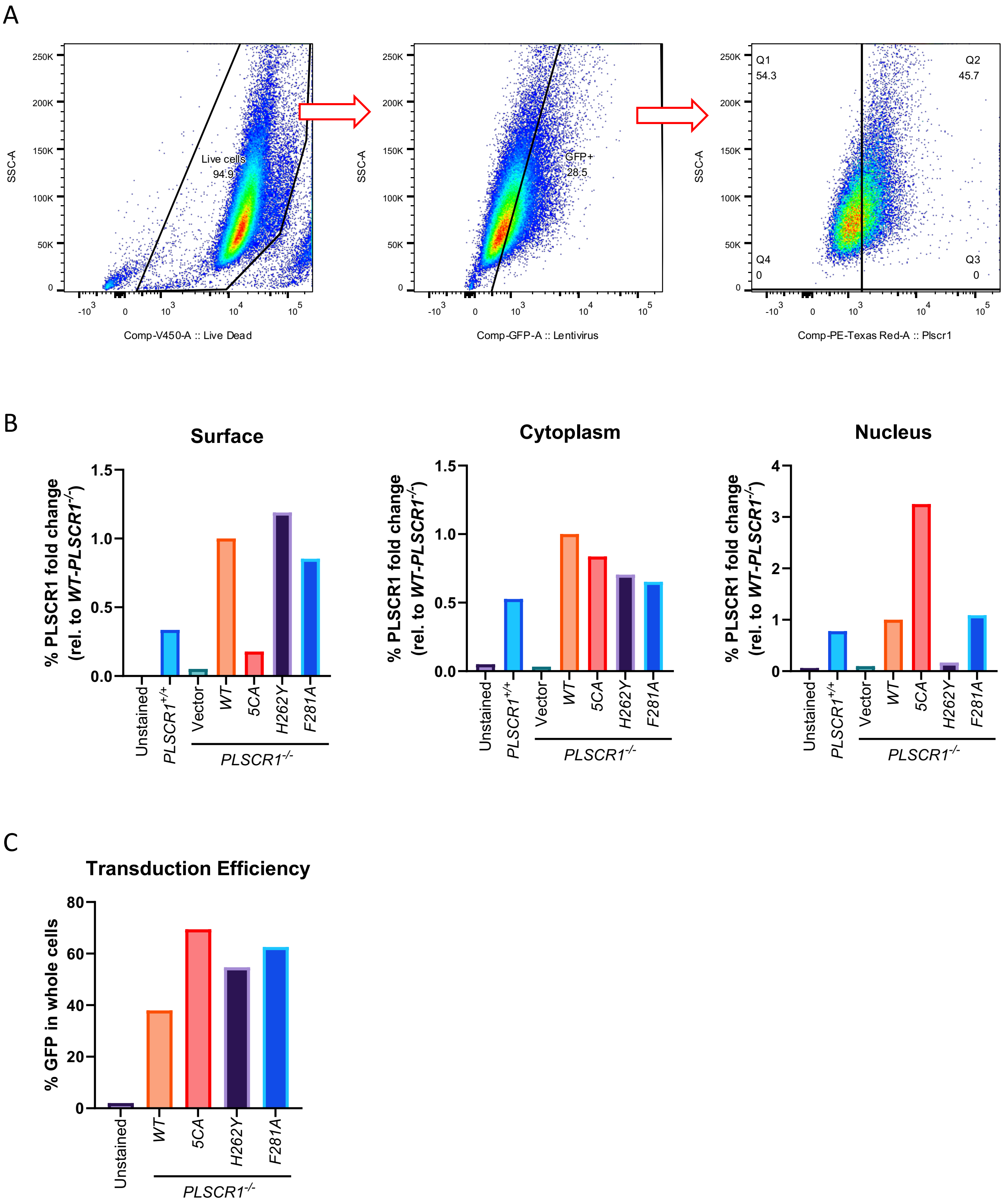


**Supplemental Figure 4. PLSCR1 Transduction Efficiency and Distribution**

*PLSCR1* plasmids on PLV-EF1a-IRES-Hygro backbone were packaged into GFP-expressing lentivirus. *PLSCR1*$^{-/-}$ A549 cells were transduced using lentivirus. After a 10-day hygromycin selection, cells were analyzed using flow cytometry.

(A) Gating strategy for live, GFP+ and PLSCR1+ A549 cells.

(B) Surface, cytoplasm and nuclear expression of PLSCR1. Data are presented as fold change compared to *WT-PLSCR1*$^{-/-}$ A549 cells.

(C) Lentiviral transduction efficiency.

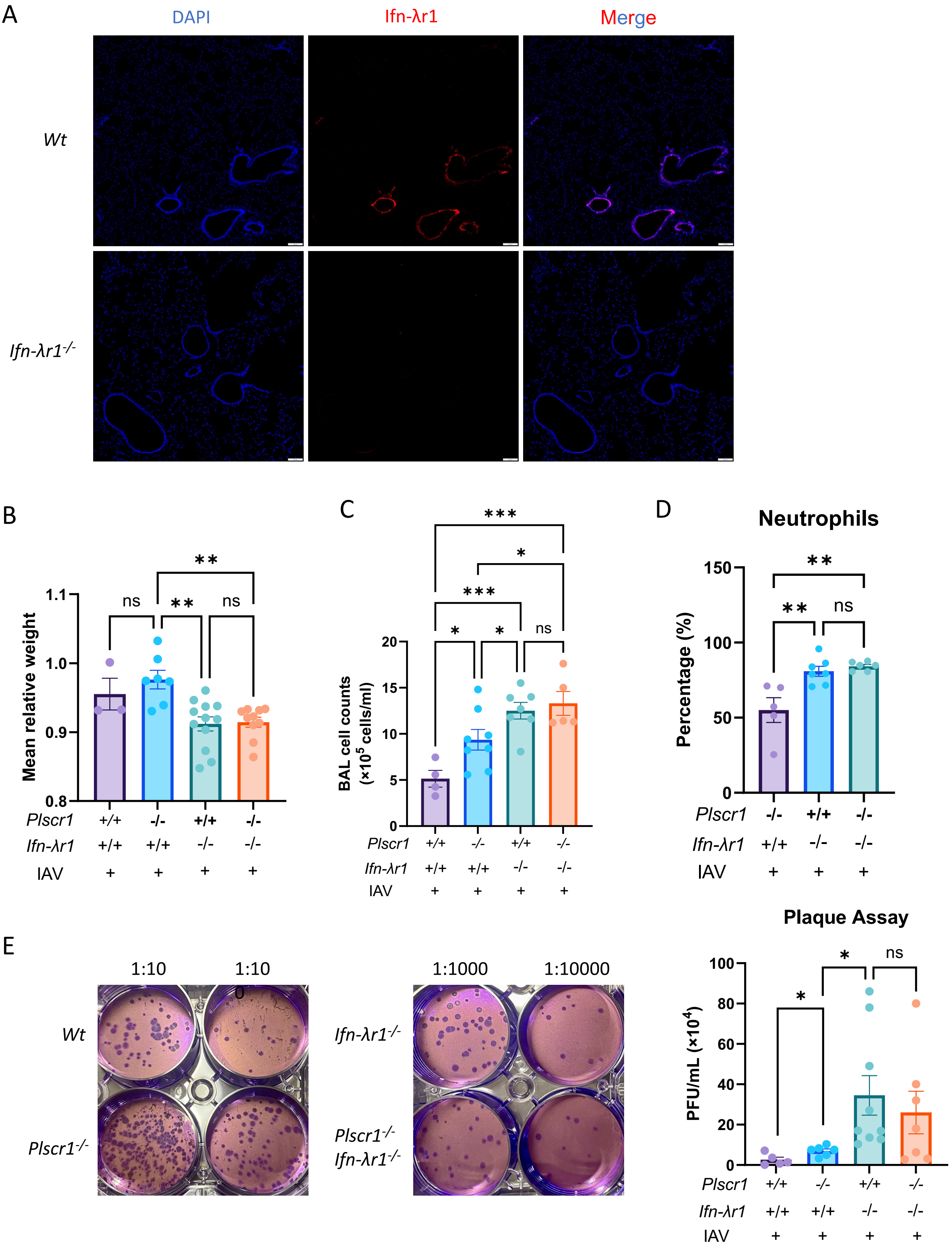


**Supplemental Figure 5. The Relative Contribution of the Type 3 IFN Pathway to Plscr1-Mediated Antiviral Immunity.**
*Wt*, *Plscr1*$^{-/-}$, *Ifn-λr1*$^{-/-}$ and *Plscr1*$^{-/-}$*Ifn-λr1*$^{-/-}$ mice were exposed to sublethal (300 pfu) IAV (WSN) infection and sacrificed at 3 dpi.
(A) Representative immunofluorescent staining for DAPI and Ifn-λr1 in lungs.
(B) Mean relative weight of mice.
(C) Total BAL leukocyte numbers.
(D) Neutrophil percentages in BAL.
(E) Infectious viral titer in the lungs was assessed by plaque assays.

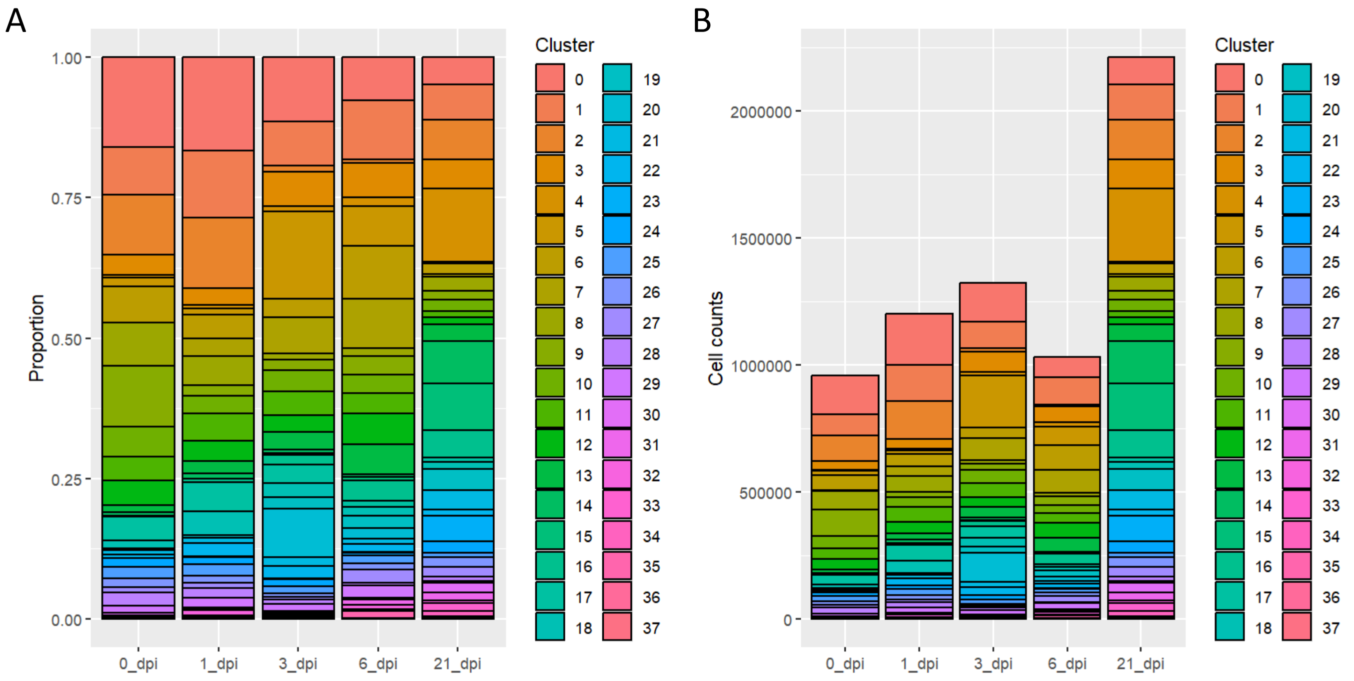


**Supplemental Figure 6. Proportion and Cell Count of Each Cluster in Single-Cell RNA Sequencing.**
*Wt* mice were exposed to 2500 EID50 IAV (PR8) infection. Lungs were used for single-cell RNA sequencing analysis at 0, 1, 3, 6 and 21 dpi.

(A) Proportion of each cluster.

(B) Cell count of each cluster.

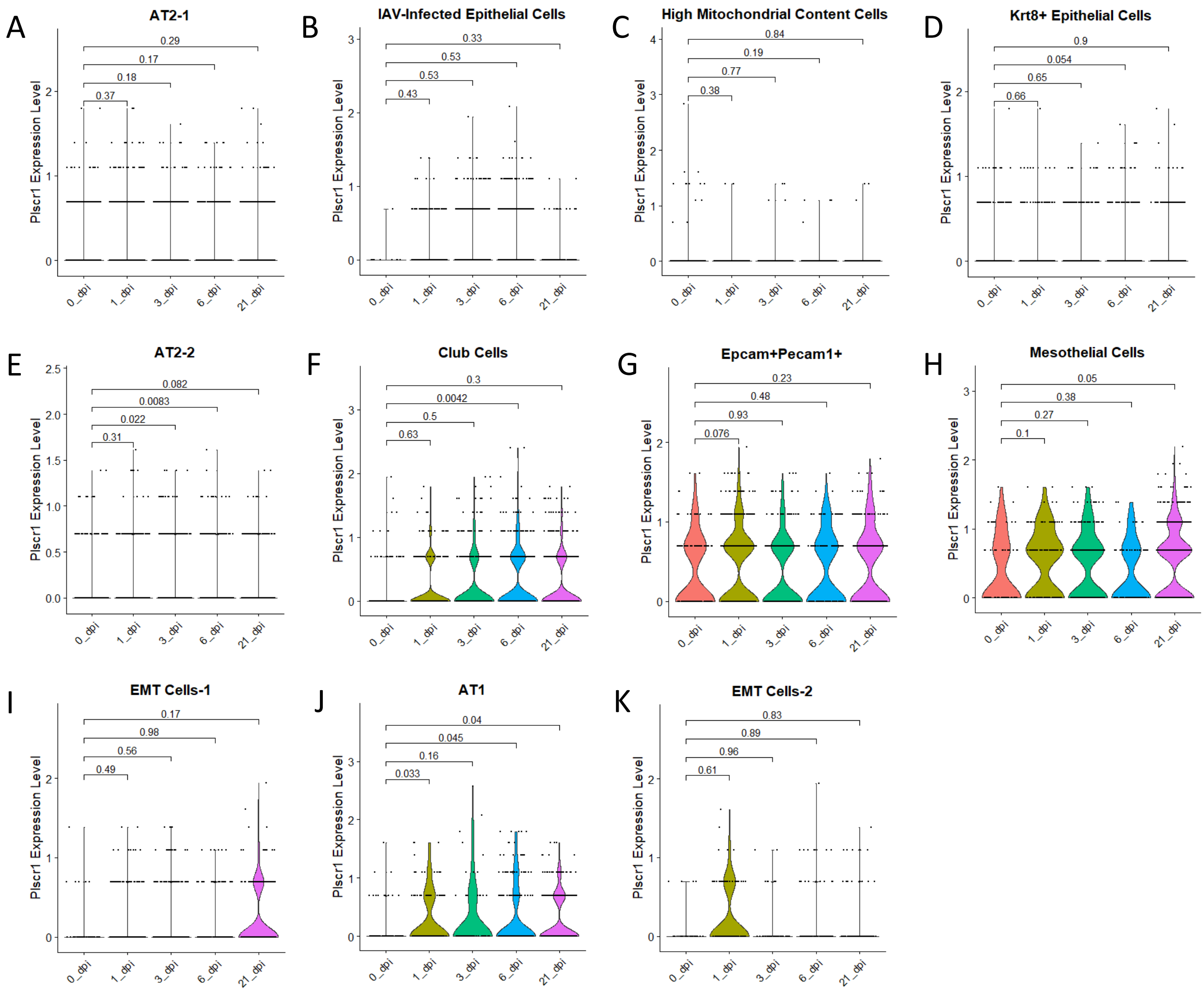


**Supplemental Figure 7. Time-Dependent *Plscr1* Expressions in All Epithelial Cell Clusters Other than Ciliated Epithelial Cells.**

*(A-K) Wt* mice were exposed to 2500 EID50 IAV (PR8) infection. Lungs were used for single-cell RNA sequencing analysis at 0, 1, 3, 6 and 21 dpi.

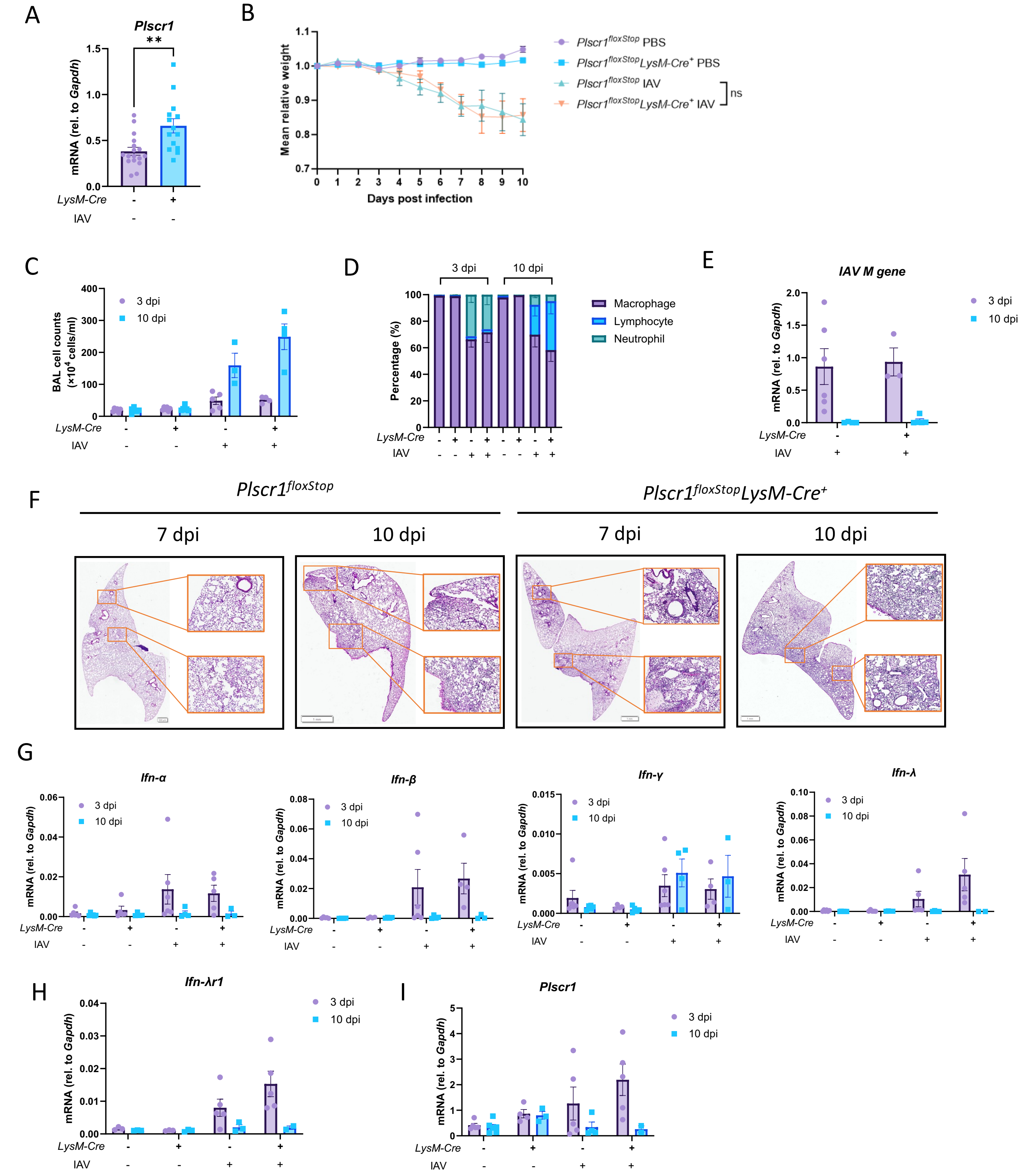


**Supplemental Figure 8. Unaffected Susceptibility of *Plscr1*$^{floxStop}$*LysM-Cre*$^{+}$ Mice to Influenza Virus Infection**

*Plscr1*$^{floxStop}$ and *Plscr1*$^{floxStop}$*LysM-Cre*$^{+}$ mice were exposed to sublethal (300 pfu) IAV (WSN) infection.

(A) Validation of *Plscr1* overexpression in lungs of *Plscr1*$^{floxStop}$*LysM-Cre*$^{+}$ mice by qRT-PCR.

(B) Mean relative weight of mice.

(C) Total BAL leukocyte numbers.

(D) Differential cell counts in BAL.

(E) Viral RNA load in the lungs was assessed by quantifying M gene by qRT-PCR.

(F) Representative lung sections stained with H&E.

(G-I) Whole lungs were analyzed for *Ifn-α*, *Ifn-β*, *Ifn-γ*, *Ifn-λ* (G); *Ifn-λr1* (H); and *Plscr1* (I) RNA by qRT-PCR.

Data are expressed as mean ± SEM of n = 15-16 mice/group for weight loss. For the rest analysis, n= 3-7 mice/group. All data were pooled from three independent experiments. ns, not significant, $*p < 0.05$, $**p < 0.01$. dpi, days post infection.

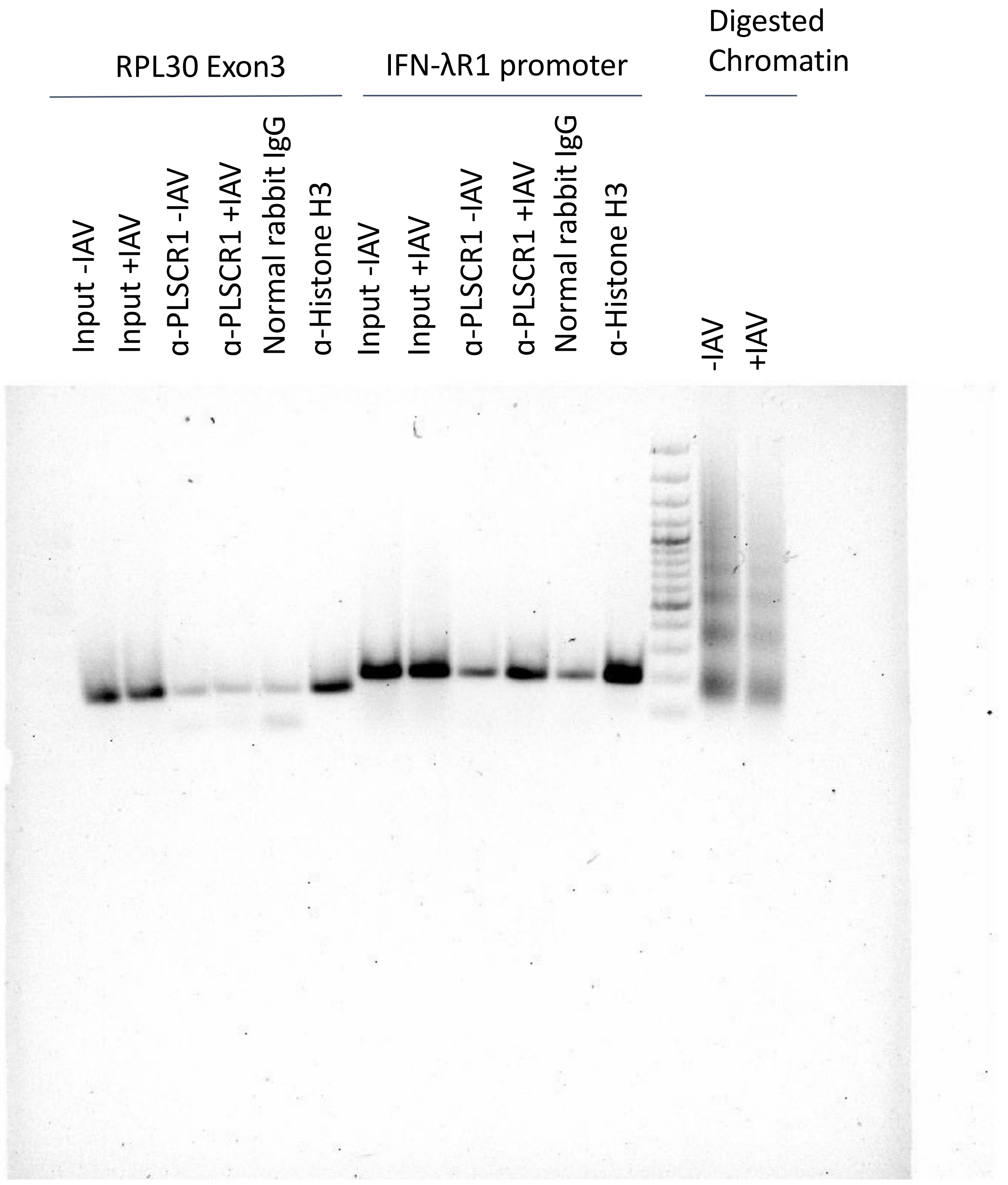
RPL30 Exon3
IFN-λR1 promoter
Digested Chromatin
Input -IAV
Input +IAV
α-PLSCR1 -IAV
α-PLSCR1 +IAV
Normal rabbit IgG
α-Histone H3
Input -IAV
Input +IAV
α-PLSCR1 -IAV
α-PLSCR1 +IAV
Normal rabbit IgG
α-Histone H3
-IAV
+IAV
